# Self organisation of invasive breast cancer driven by the interplay of active and passive nematic dynamics

**DOI:** 10.1101/2024.12.09.626295

**Authors:** Pablo Gottheil, Saraswat Bhattacharyya, Kolya Lettl, Philip Friedrich, Kilian Roth, Salvador Rivera-Moreno, Mario Merkel, Bahriye Aktas, Igor Sauer, Assal Daneshgar, Jonas Wieland, Hans Kubitschke, Anne-Sophie Wegscheider, Julia M. Yeomans, Josef A. Käs

## Abstract

In invasive breast cancer, cell clusters of varying sizes and shapes are embedded in the fibrous extracellular matrix (ECM). Although the prevailing view attributes this structure to increasing disorder resulting from loss of function and dedifferentiation, our findings reveal that it arises through a process of active self-organization driven by cancer cell motility. Simulations and histological analyses of tumours from over 2,000 breast cancer patients reveal that motile, aligned cancer cells within clusters move as active nematic aggregates through the surrounding highly aligned ECM fibres, which form a confining, passive nematic phase. Cellular motion leads to cluster splitting and coalescence. The degree of cluster activity, combined with heterogeneity in cell motility, is reflected in specific scaling behaviours for cluster shape, size distribution, and the distance between cluster boundaries and nematic defects in ECM alignment. Increased activity estimates correlate with tumour progression and are associated with a poorer prognosis for patients.

---

Breast cancer starts as an uncontrolled growing, non-invasive cell mass encapsulated by extracellular matrix (ECM), called a tumour *in situ* [1] (Fig. 1b). Progressed invasive lesions show a more complex structure: many smaller, heterogeneous clusters of cancer cells are embedded into dense, fibrotic ECM (Fig. 1d) produced by cancer-associated fibroblasts [2]. We have recently shown that cells within a cluster can undergo a collective cell motility transition [3] in addition to the processes of cell proliferation and ECM production. However, how self organisation dynamically generates the heterogeneous and complex tumour structure, which serves as a clinical signature for the onset of malignancy [4], has remained unclear.

**Fig. 1.**
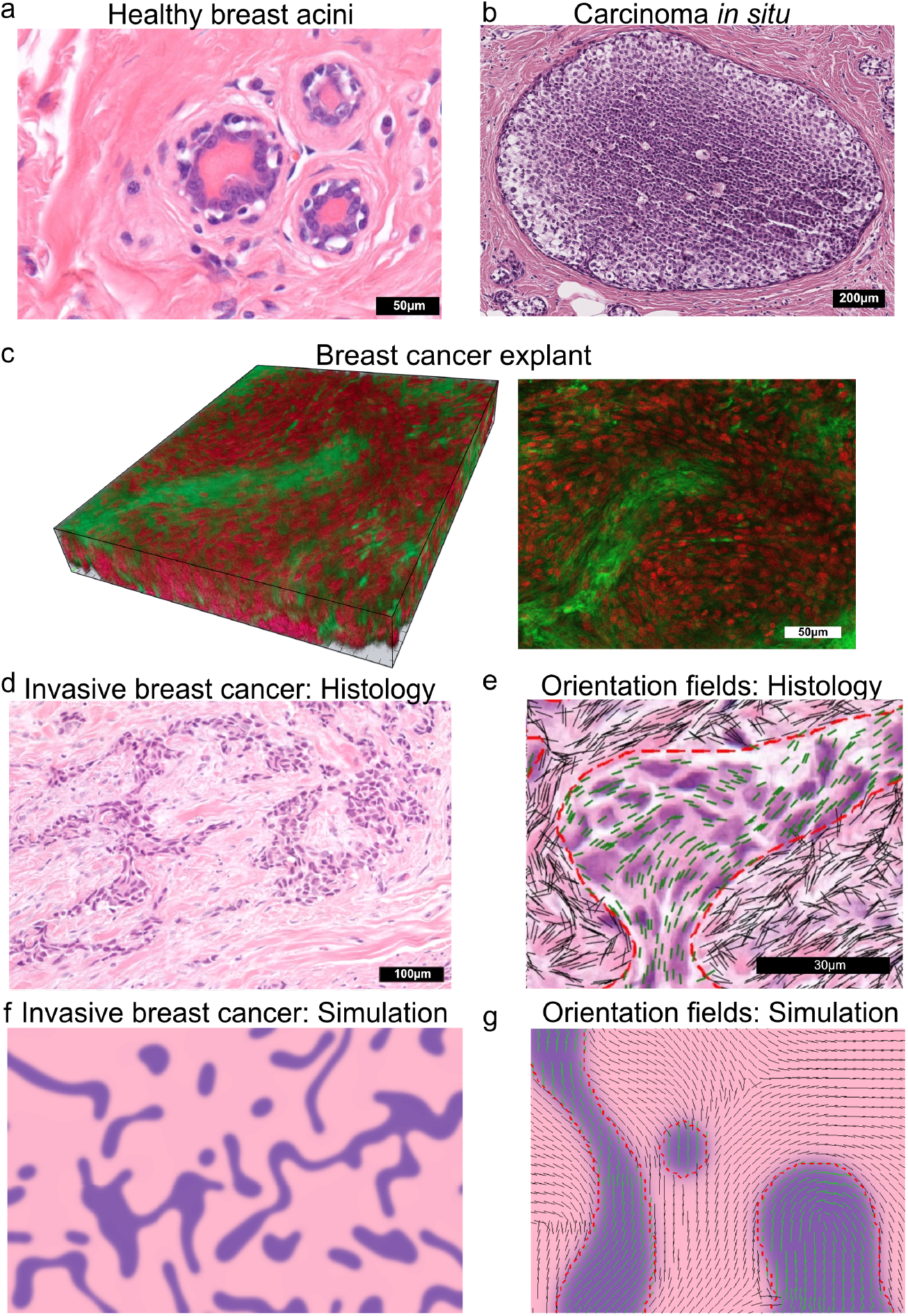
Structure of solid breast tumours. H&E staining shows cell nuclei in purple and extra-cellular matrix (ECM) in pink. **a** Healthy breast epithelium confined by ECM fibres forming a basal membrane. **b** The ductal carcinoma in situ (DCIS) is an early in situ tumour defined by uncontrolled growth of neoplastic epithelial cells only in the (milk) duct, still confined by an intact basal membrane **c** 3D volume of primary breast cancer (316*µm* × 253*µm* × 40*µm*) stained for DNA (red) and actin (green) exhibits 3D cell nematic order (Supplementary Note 1.1.2). **d, e** Later stage invasive lesions show a more complex structure: **d** native histological image, **e** orientation fields. Many smaller, highly irregular cancer cell clusters with well-defined boundaries are embedded in the collagen fibres of the ECM. **f, g** The assembly of this heterogenous tumour structure can be understood by simulations that rely on liquid crystal physics: **f** simulation phase field, **g** orientation fields. In the clusters the cells form aligned streams of moving cells (green lines in **e, g**). The surrounding ECM consists of regions of collagen fibres with local alignment (black lines in **e, g**).

We compare extensive histological images of breast cancer patients and continuum modelling to show that the mechanical interplay that leads to early invasive tumour lesions can be understood using concepts from liquid crystal physics. In particular, the local alignment of ECM fibres is characteristic of nematic ordering, where the term nematic denotes a preferred alignment axis [5]. Moreover, recent work has shown that cellular unjamming can lead to streams of cells in the clusters [6, 7]. The collectively moving cells can be described as an active nematic, where *active* indicates that the cells are motile [8, 9]. As the clusters move, active stresses across their boundaries which result from cell motion lead to a dynamical steady state of clusters deforming, breaking up and merging. Hence the distinctive pattern of invasive lesions highlights the role of collective cell motility.

## Histological images reveal nematic order in cancer aggregates and ECM

To comprehensively assess the range of tissue structures found in tumours we analysed digital histological slides of 2170 breast cancer patients (*N* ≈ 1.8 · 10^5^ clusters). Such a large patient cohort guarantees reliable data on cancer cluster size distribution and shape, as well as nematic properties of the cells and ECM, rendering it an ideal starting point for the comparison to our continuum modelling. We also included 87 patients with ductal *in situ* carcinoma from an open histology database [10] (*N* ≈ 5 · 10^2^ clusters), and 32 samples from healthy tissue regions that serve as a reference.

The image analysis of H&E-stained tumour cutouts (see Methods) measures the nematic order of the ECM based on the eosin-stained fibres, and the nematic order of the cell clusters based on hematoxylin-stained nuclei. The fibres of the ECM are clearly aligned forming polymeric, passive nematic domains with an average order parameter of 0.52 ± 0.07. Clusters contain motile cells [3, 11] that align to an active nematic phase (compare Figs. 1c-e) with an average nematic order parameter of 0.46 ± 0.22 in the histological sections. Dimensionality analyses of cell and ECM order are given in Supplementary Note 1.1. As visible in Figs. 1b,d-e the cancer cell clusters display a sharp boundary indicating that cell-cell attraction prevails over cell-ECM adhesion. This generates surface tension between cell clusters and ECM. Nevertheless, the clusters have a non-spherical, irregular shape, which suggests that mechanical stress which results from the cell motility deforms the cluster shape and exerts forces on the ECM.

## Invasive cancers can be described as active nematic materials

These findings motivate the hypothesis that the collective behaviour of invasive lesions can be described as active nematic inclusions which exert deforming stresses on a surrounding viscoelastic nematic phase of ECM fibres. However, each histological image corresponds to a fixed time. To gain insight into the dynamical mechanisms controlling the distribution and evolution of clusters in tumours, we simulate a viscoelastic nematic phase, representing the ECM, coupled to an active nematic phase, representing the cancer cell clusters (described in Methods).

Mapping from simulation to physical parameters follows from the choice of a length, time and stress scale. Typical cluster velocities based on our measurements of cancer cell motility [3] are ∼3*µ*m h^−1^ and typical cell radii ∼5*µ*m. The typical bulk modulus of a breast tumour sample, that is dominated by ECM, is ∼500Pa as our AFM measurements of primary human breast cancer (see Supplementary Note 1.2 and Methods) and prior publications [12, 13] confirm. Traction force microscopy shows that the cancer cells typically exercise mechanical stresses of about a few 100 Pa on the ECM [14, 15]. These experimental values guided our choice of parameter values used in the simulations. Details of the mapping and a discussion of the robustness of the model are given in Supplementary Note 1.3.

The 2D simulations show a large number of mobile, tortuous cancer cell clusters (Fig. 1f) that are propelled and stretched by active flows produced by the motile cells. The mechanical forces due to activity are able to extend the clusters and overcome surface tension, so that the clusters break up, move around, collide and re-form [16–18]. This dynamical evolution is illustrated by Supplementary Movie 1 and the time sequence in Fig. 2a. Snapshots of the cluster distribution for two different activities are compared to histological data in Figs. 2b. Fig. 2c collates simulation data on the variation of cluster size with activity. These figures show that the cluster size is reduced by higher activities which foster the clusters to extend and break up the cell aggregates. To check whether we were considering a dynamical steady state independent of initial conditions, we ran the model starting from both a single large tumour or a large number of tiny tumour clusters, with the cells occupying an area fraction *ϕ* = 25%. In both cases, the clusters reached the same steady-state distribution. The empirically observed area fraction is 0.41 ± 0.17 (patient median ± median absolute deviation). We check the influence of *ϕ* on the simulation results in Supplementary Note 1.3.

**Fig. 2.**
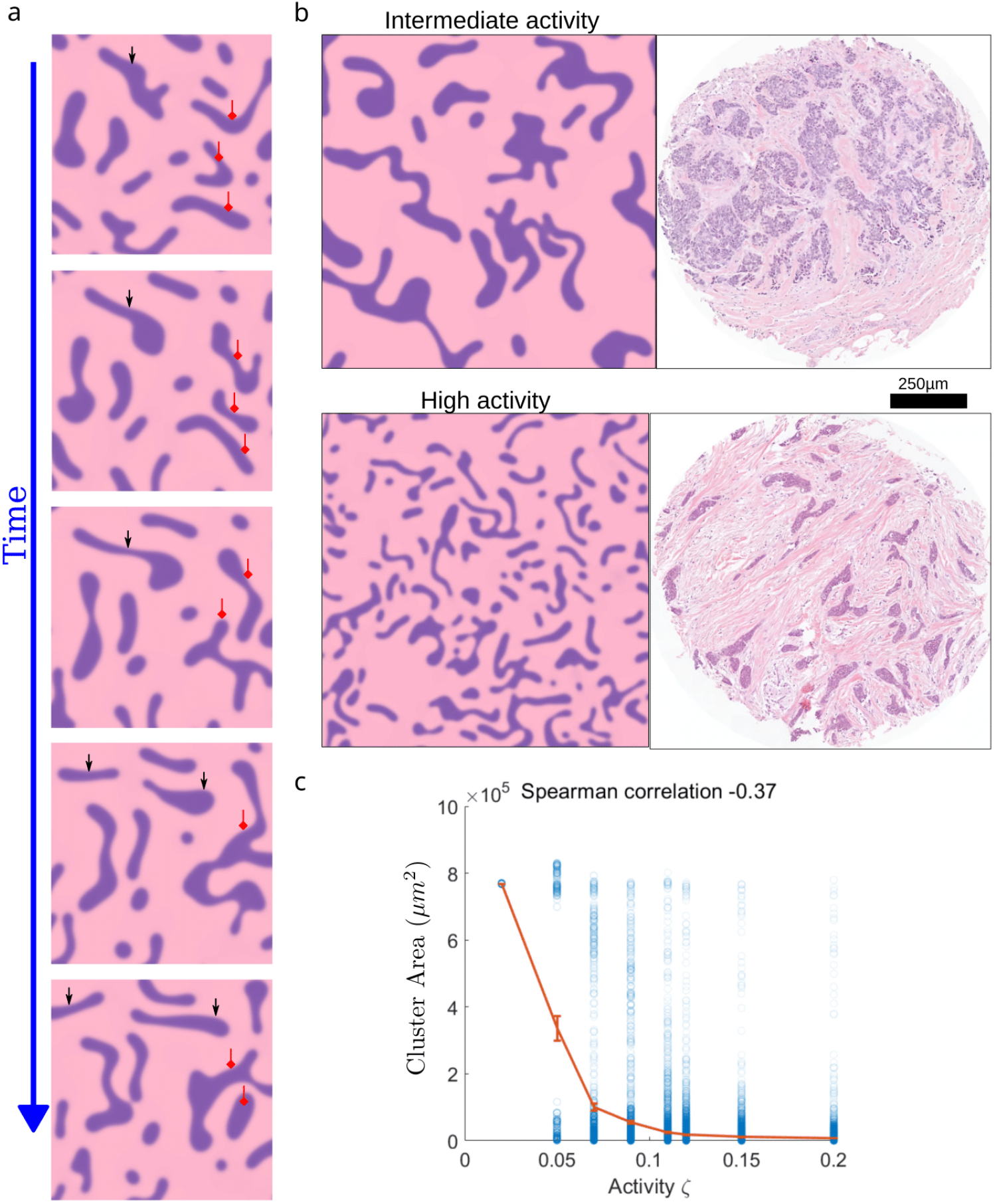
Mechanical activity in the cancer cell clusters caused by cell motility determines the shape and size of the clusters. **a** Time sequence from simulations showing the cluster dynamics, with droplets moving, splitting, and merging. Black (red) arrows indicate examples of clusters breaking up (recombining) over time. **b** With increasing activity the clusters become smaller. Simulations are compared to examples from the histological images chosen to show similar structures. **c** Measured cluster sizes as a function of activity from simulations (N=44,714 clusters). The cluster sizes show a negative correlation with activity.

## Cancer cluster area and shape distributions

These results were unexpected as current tumour biology only reports that motility facilitates cancer cell escape from the clusters into the ECM by single cells or small cell aggregates breaking away [7] rather than dynamics where cluster fission and fusion impact the size distribution. Therefore we next obtained the cluster area and shape distributions from the histological images and compared these to the distributions from our simulations. According to our previous experiments [3, 6] clusters in a tumour vary in the fraction of motile cells that they contain due to inherent tumour heterogeneity. To reflect this we assigned the clusters a range of different activities in our simulations.

The probability distribution of cancer cluster areas in histological images of invasive breast cancer as a measure of cluster size (*N* ∼ 10^5^ clusters) is plotted on a log-log scale in Fig. 3a (red line), showing a clear power law distribution of cluster sizes *P* (*A*) ∼*A*^*α*^ with *α* =− 2.38 with 95% confidence interval [ −2.43, −2.33] (*R*^2^ *>* 0.99) over a range of areas from ∼3· 10^2^*µ*m^2^ −3· 10^5^*µ*m^2^ with an average value ⟨*A*⟩ = 3.7 ·10^3^*µ*m^2^. The simulations reveal the same power law exponent from ∼10^3^*µ*m^2^ −10^5^*µ*m^2^ (*R*^2^ = 0.98, Fig. 3a, blue lines). The first three bins from the simulations do not exhibit the power law scaling because of the difficulty of resolving the smallest clusters in the simulation phase field. We note that the images represent 2D cross sections through 3D tumours, and the simulations were carried out in 2D, this point is discussed in Supplementary Note 1.1.3. Interestingly, 3D volumes of pancreatic intraepithelial neoplasia aggregates - a precursor for invasive ductal adenocarcinoma, were also found to be power-law distributed [19].

**Fig. 3.**
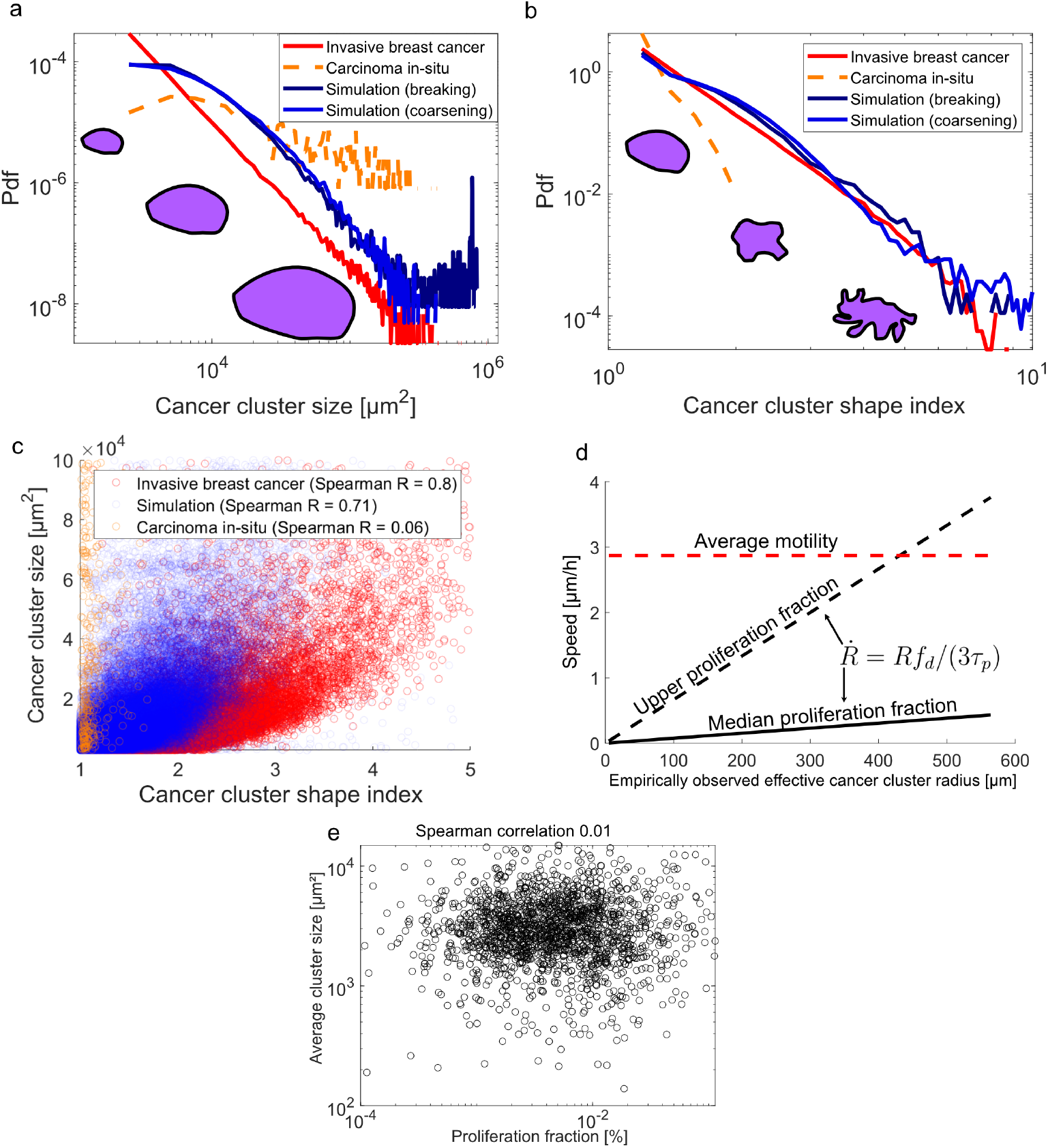
Scaling of shape and size in breast tumour cell clusters. Distribution of **a** cluster sizes. **b** cluster shapes 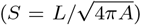. In each graph, the red line displays the result from analyzing the histological slides from 2012 patients. Data from our simulations, with an initial condition of one large tumour cluster or many tiny clusters, are shown in deep blue and blue respectively. The yellow dashed line shows results for *in situ* breast tumours that do not show a scaling behaviour. Inset cartoons give a pictorial guide to changing sizes and shapes. **c** The size and shape of invasive lesions are strongly correlated in both histological data from breast cancer patients (blue symbols; Spearman *R* = 0.80) and simulations (red symbols; Spearman *R* = 0.71 for clusters *>* 2000*µm*^2^). There is no correlation for *in situ* breast tumours (yellow symbols; Spearman *R* = 0.06). **d** Movement of the clusters can be caused by cell motility or proliferation: Only in very large clusters can proliferation dominate, in 99% of all clusters cell motility is the dominant factor. **e** There is no correlation between cluster size and proliferation fraction. This is consistent with the hypothesis that cell motility is responsible for cluster size distribution.

On increasing activity, not only the size but also the shape of the cluster changes. To quantify the irregular shapes, we measured the perimeter *L* of each cluster within the ECM and defined the cluster shape index 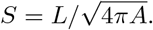 is 1 for a circular cluster and increases from unity as the boundary of the cluster becomes more tortuous. The probability distribution of *S* for the histological images is shown in Fig. 3b (red line), together with the distribution of *S* for simulated clusters (blue lines). The mean value of *S* of the invasive breast cancer clusters is 1.38. Due to the small range of *S*, we are unable to distinguish between a power law distribution (∼ *S*^*β*^; *β* = −5.65 ± 0.29 (fit ± 95% CI), 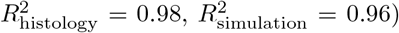 and an exponential distribution 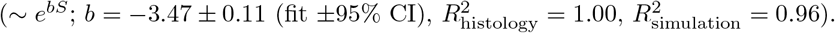

The size and shape distributions are closely related to each other, as can be seen by the direct correlation of the two quantities (Fig. 3c). The corresponding Spearman coefficient for invasive breast cancer clusters is *R* = 0.8 and for simulations *R* = 0.71, reflecting strong positive correlations between size and shape – clusters with larger area are more tortuous in shape. This close agreement between data and simulations gives further confidence in the physical interpretation of the distribution of the cancer clusters as a dynamical steady state arising from a balance between active cluster break up and fusion.

As a comparison, we analysed the shape and size distributions of clusters of ductal *in situ* carcinomas. This led to the yellow, dashed curves in Figs. 3a-c. The most striking difference to invasive tumours is the round regular shape of the cancer cell aggregates. The cluster shape index never exceeds 2 and over 80% of clusters exhibit indices of less than 1.2 (dashed line, Fig. 3b). There is no correlation between cluster shapes and sizes (Fig. 3c). Moreover, the distribution of cluster sizes is weighted away from small sizes towards larger sizes when compared to the later invasive lesions (Fig. 3a). This confirms the difference between *in situ* tumours which are just a growing cell mass confined by the basal membrane and the highly motile, later stage, invasive clusters.

In growing tumours, cell division can result in active flows of a similar nature to cell motility. We now estimate the relative contributions of proliferation and motility to the activity of the clusters in the invasive lesions. Assuming spherical clusters of radius *R*, the average spreading speed caused by cell proliferation is 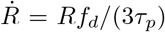 where *f*_*d*_ is the fraction of mitotic, i.e. actively dividing, cells and *τ*_*p*_ is the typical duration of mitosis. We assumed *τ*_*p*∼_ 2h [20] and measured the fraction of mitotic cells in the patient collective using the proliferation marker Phospho-histone H3 (see Methods), giving a median value of *f*_*d*_ ∼0.46% and an extreme value of ∼4% for the 99th percentile of all patients. The variation of proliferation-induced speed with cluster radius for observed radii and the median and extreme estimates of *f*_*d*_ are shown in Fig. 3d where they are compared to an estimate of the average speeds of unjammed cells inside breast tumour explants, ∼3*µ*m/h, obtained from our previous vital cell tracking experiments [3]. Although these are rough estimates, they provide evidence that motility dominates proliferation in driving the motion of almost all cancer clusters. The lack of any correlation between cluster size and proliferation fraction (Fig. 3e) further suggests that motility, instead of growth, is responsible for the observed distribution of cluster sizes.

## Activity leads to cluster break up and topological defects in the surrounding nematic ECM

To investigate further how the clusters break up dynamically, we performed simulations starting from a single cluster and varying the range of activities and the mechanical resistance of the ECM within the experimentally observed values. The resulting state diagram is shown in Fig. 4. To the left of the red dashed line, for smaller activities and larger ECM resistance, the cluster remains intact, but as the border of this region is approached, active forces distort the cluster to a fluctuating, and on average more elongated, shape. To the right of the red dashed line, the higher activities are able to overcome surface tension and pull the tumour apart into a large number of motile smaller clusters (see Fig. 2a). For exemplary points in the state diagram, in the stable and unstable regions, we display a typical cell cluster together with a histological slide to show that the different cluster shapes are observed in real tumour tissue observed in histopathological diagnostics. The dynamics of these representative clusters are shown in Supplementary Movies 3a–3e.

**Fig. 4.**
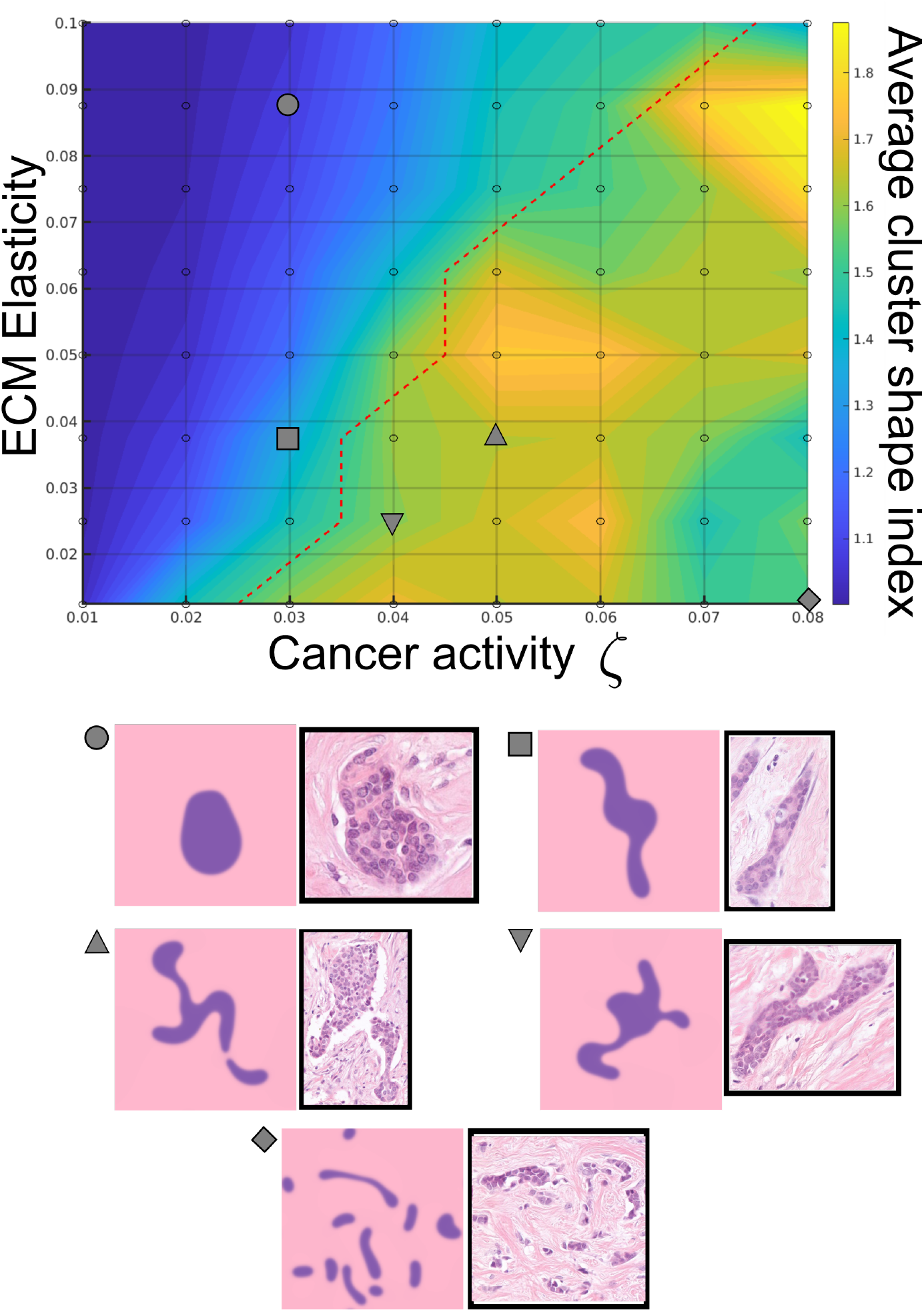
Stability of tumour cell clusters. The state diagram shows the cluster shape index (heat map) as a function of the mechanical resistance of the ECM and the activity of the cluster. To the right of the red dashed line, a single active cell cluster breaks apart into multiple clusters. For the points marked by symbols in the diagram, we compare typical cluster shapes from simulations and histological images.

We next investigate how moving clusters interact with the surrounding ECM (Fig. 5a). Previous literature predominately described the mechanical interactions between cell clusters and ECM in terms of the traction forces that individual cancer cells or small aggregates generate when trying to escape the clusters [14, 21, 22]. One prevailing opinion is that the traction forces pull on the ECM and align it radially with respect to the cluster boundary, which fosters cell escape [23–25] and lower tissue surface tension. While it was experimentally shown that single-cell escape is a valid escape mechanism [7], our modelling suggests that the collective motion of the active clusters has a significant effect on the surrounding ECM.

**Fig. 5.**
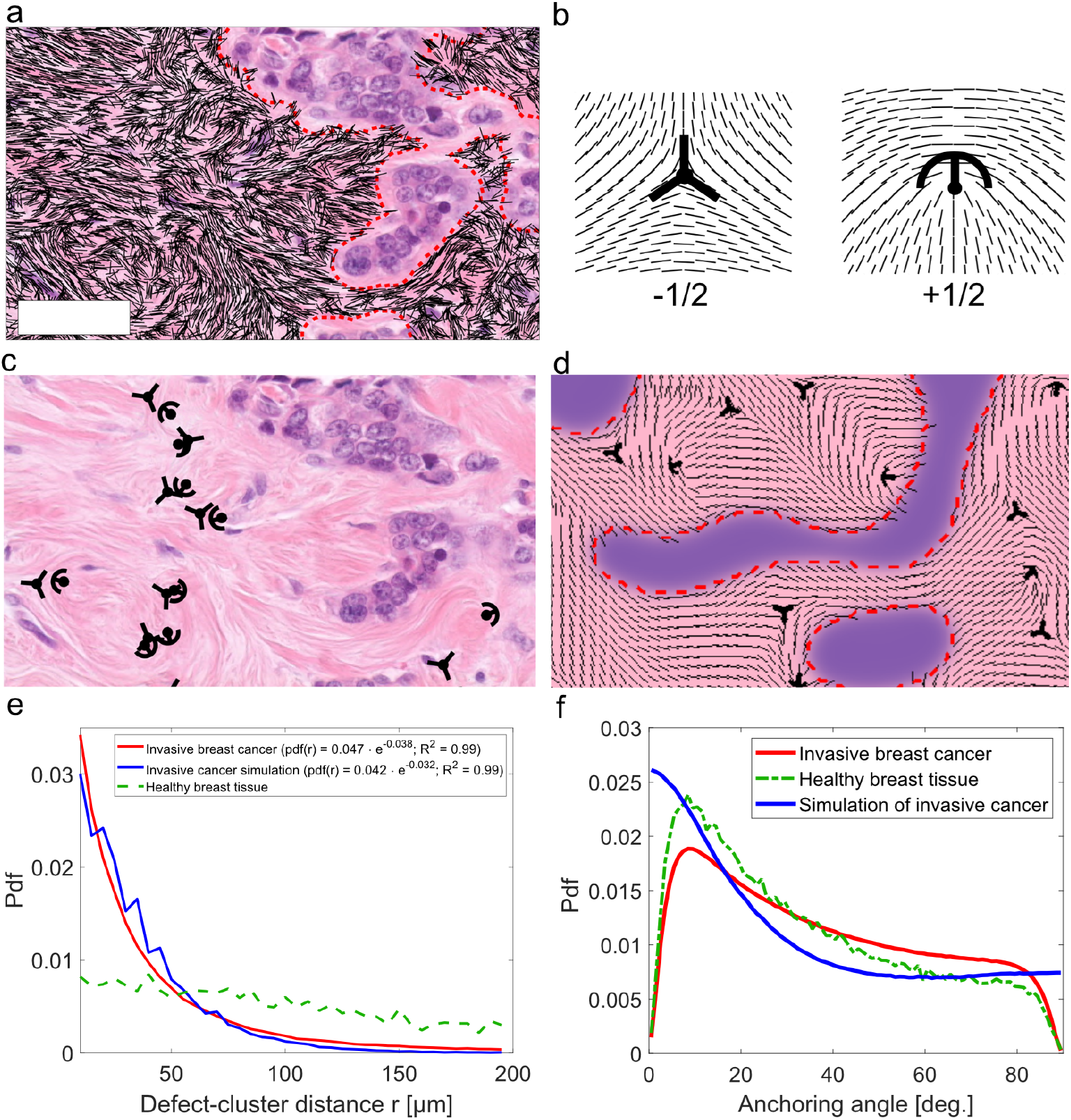
Effects of activity on the ECM. **a** Nematic ordering in the ECM, shown by the black lines. The scale bar is 100*µ*m. **b** Alignment around a −1/2 (trefoil) and +1/2 (comet) topological defect. **c**,**d** Topological defects in the ECM surrounding breast cancer lesions in c) a histological slide and d) a simulation snapshot. **e** Distribution of the shortest distance between defects in the ECM and a cluster boundary. The number of defects decreases exponentially with the distance from the cluster boundaries in invasive breast cancer tissues (red) and active nematic simulations (blue) but is relatively constant in healthy tissues (green). **f** The nematic ECM predominantly aligns parallel to the active cluster boundary.

We compared the nematic ordering of the fibres within the ECM between healthy breast tissue and the invasive lesions (see Supplementary Note 1.4). While both show a wide variability, the invasive lesions do not show higher nematic order parameters despite that cancer-associated fibroblasts generate additional fibrotic ECM in the tumour which leads to an increase in density [26].

Local disruptions in the alignment of the nematic ECM generate topological defects, points where the ordering of the ECM locally takes the form of a comet (+1/2 defect) or trefoil (−1/2 defect) (Fig. 5b). Figs. 5c,d provide visual examples of defects in the director field for the histological images and simulations respectively. These correspond to disclinations in the 3D ECM (Supplementary Note 1.1.4). The defects tend to be formed in the vicinity of the tumour cell clusters (Fig. 5e), and the number of defects decreases exponentially with the distance from the cluster boundaries in both histology and simulations. However, defects in healthy tissue are more uniformly distributed throughout the ECM. This suggests that the defects surrounding the invasive lesions are formed due to stresses caused by the activity of the tumour cell clusters.

We then compared the orientation of the nematic director of the ECM at the cluster boundary in the histological sides with simulations. In both clinical data and simulations, the director preferentially aligns parallel to the boundaries of the tumour cell clusters. This is quantified in Fig. 5f which plots the angle between the tangent to the boundary and the nematic direction, showing a clear maximum at ∼0^°^ and qualitative agreement of the simulations with the data. This suggests that the ECM aligns to the flow field produced by the active clusters. Anchoring dimensionality is discussed in Supplementary Note 1.1.5.

## Activity influences tumour progression

Active forces can explain the self-organisation that leads to the characteristic structure found in invasive breast cancer lesions that comprise small islands of motile cancer cells embedded into passive ECM. However, the important question remains of whether the self-organisation process is essential for tumour progression.

As indicators of cluster activity that drives the formation of invasive lesions, we used the cluster shape index (see Fig. 3b), and the normalized maximum defect-cluster distance (detailed in Methods). We applied the Kaplan-Meier estimator [27] to determine whether the survival function depended on each of these parameters. The patient collective was divided into a training group (30% of all patients) to find the optimal threshold in our activity estimates to distinguish between low and high risk and a test group (remaining 70% of all patients) to validate our prognostic stratification criteria. Details of the parameters used for risk assessment and thresholds can be found in the Methods. Complete information on the patient collectives used for the retrospective study can be found in Supplementary Note 1.5.

The Kaplan-Meier plots (Fig. 6a) show that a higher average cluster shape index correlates with a worsened prognosis. In our model, a higher shape index corresponds to active droplet break-up, corresponding to higher activities and lower elasticity near cancer clusters (Fig. 4). The next Kaplan-Meier plots (Fig. 6b) show that a low defect-cluster distance also corresponds to a worsened prognosis. We interpret this as an interplay of activity and elasticity, where topological defects created by active forces remain close to the cancer cluster due to the elasticity of the ECM.

**Fig. 6.**
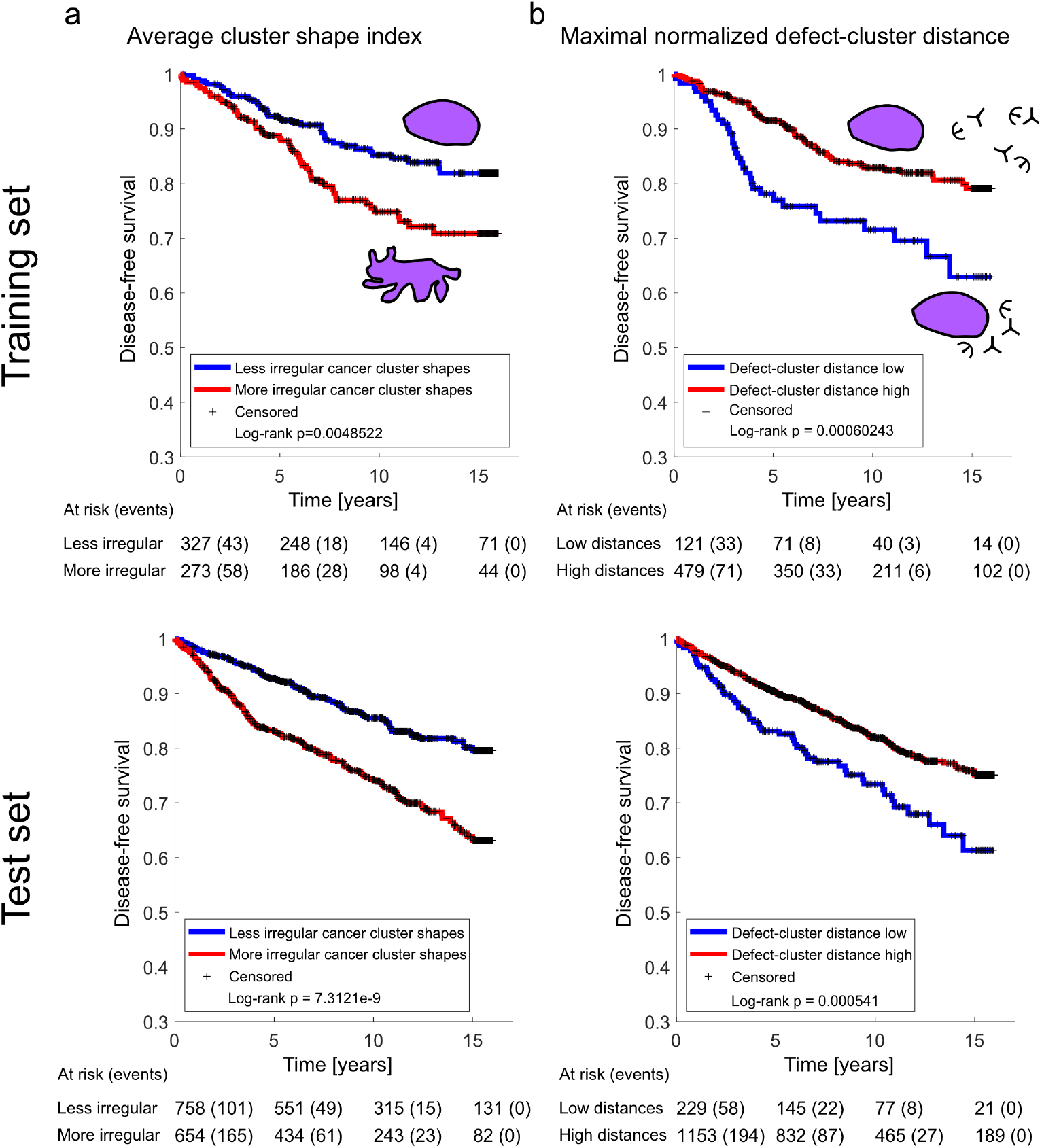
Survival curves are shown for a random 30% training and 70% test set (detailed patient statistics in Supplementary Note 1.5). In the training set, we find the threshold that maximizes risk group separation according to the log-rank statistic. The threshold is subsequently applied to the test set patient collective. Disease-free survival probabilities are plotted according to Kaplan-Meier estimators. **a** The average cluster shape index was used for risk group separation in training (N = 600) and test (N = 1412) collectives. The log-rank p-value in the training set is *p ≈* 0.005 and in the test set *p <* 0.001, respectively. **b** The maximum defect-cancer distance normalized to the ECM length scale was used for risk group separation in training (N = 600) and test (N=1382) collectives. The log-rank p-value in the training set is *p <* 0.001 and in the test set *p <* 0.001, respectively. *At risk* indicates the total number of patients who have not been censored, developed distant metastasis or died by a specific follow-up time. In parentheses after the *At risk* is the cumulative number of individuals that developed distant metastases or died later than indicated.

These results suggest that it would be interesting to investigate further the role of active mechanical forces in cancer progression. The process is, however, likely to be complex: cluster instabilities and breakage increase the total surface between cell clusters and ECM. Small clusters foster exchange with the microenvironment, affecting metabolic supply, chemical signalling, and mechanotransductive signalling. A recent publication demonstrates that cluster boundaries increase the selective advantage of new and more aggressive cancer cell phenotypes generated by genetic instability at the boundaries compared to new cell phenotypes emerging in the bulk of the cell clusters [28]. Moreover, the cells at the boundary are in contact with the ECM that acts as a tumour promoter [29]. Conversely, the ECM structure around the inclusions hinders cancer cell escape from the clusters into the ECM, since most ECM fibres are aligned parallel to the boundary, and constrain the cancer cells [30].

In conclusion, the pattern of tumour cell clusters within the ECM that characterises invasive breast cancer is the result of a dynamical process whereby clusters break-up, move, collide and re-form, driven by the motility of the cancer cells. These patterns can already be found in early invasive lesions after the cancer cells have broken through the basal membrane constraining a tumour *in situ* indicating an early onset of cell motility in breast cancer. Mechanically, we expect that regions of the state diagram where clusters are stable are associated with low risk patients and the regions of unstable clusters come with a high risk prognosis. This mechanical activity can be understood by describing the clusters as active nematic inclusions embedded into the ECM modelled as a viscoelastic nematic matrix. Our results suggest new experiments and may have an impact on prognostic tumour markers and inspire novel therapeutic approaches.

## Methods

### Histological tissue sections

Imaging and staining of patient breast cancer tissue was performed following standard protocols. H&E-staining was carried out using the Leica ST4040 automat. The formalin-fixated breast cancer tissue sections were then optically inspected and digitized using the Panoramic 1000 Flash IV Scanner of 3DHISTECH with a Zeiss Plan Apochromat 40x objective. Each patient is represented by a tissue micro-array core of diameter ∼1000*µ*m which was sampled from within the centre of the primary tumour. Patients were informed and asked for ethical, informed consent, approved by the ethics committee of the Ä rztekammer Hamburg (’Medical Council Hamburg’) with processing number PV2946.

### Nematic image analysis

Following a custom colour-deconvolution algorithm based on the work of Macenko et al. [31], we use the eosin-intensities to characterize ECM ordering and hematoxylin-intensities for nuclei, respectively. Cancer cell clusters are identified via an algorithm described in our previous work [3]. Using this intensity image *I*, we compute the structure tensor *J* by convolving (denoted by ∗) with a Gaussian kernel *G*(*σ*) [32],

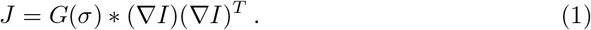

The kernel size is 2· 3*σ*, ignoring low-weight contributions greater or lower than +3*σ* and −3*σ*, respectively. We use *σ*≈ 1*µm* for the ECM fibers, and *σ* ≈5*µm* for the cancer cells.

For each pixel **p**, we need to calculate the local ECM or cell orientation **n** = (cos *θ*, sin *θ*). We do this by solving the eigenvalue problem **J**(**p**)**v** = *λ***v**. The eigenvector **v**_*min*_ corresponding to the smaller eigenvalue *λ*_*min*_ describes the local ECM or cell orientation **n** at that pixel [33]. We ignore regions where the orientation field has low coherency 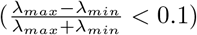 or low energy (| *λ*_*max*_ |+ |*λ*_*min*_ |*<* 0.01).

We calculate the coarse-grained nematic tensor **Q** from the orientation field **n** as

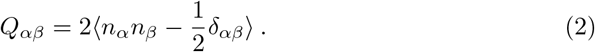

Here, δ_*αβ*_ is the Kronecker-Delta, and ⟨…⟩ denotes taking a Gaussian average over space with standard deviation *σ*. Here, we use a standard deviation of 5*µ*m for ECM fibres and 25*µ*m for cancer cells.

We find topological defects by evaluating the winding number

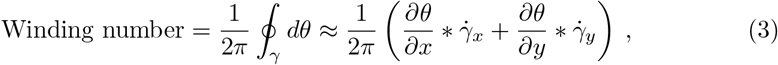

where we integrate over paths *γ* which are rings with a radius of 4 pixels, and 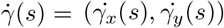 is the derivative of the path *γ*(*s*) with respect to the parametrization *s*.

### Fixation, Staining, and Fluorescence Imaging of 3D Primary Breast Tumour Sample

In Supplementary Note 1.1.2, we present findings on the relationship between 2D and 3D cell nematic ordering based on nuclei and actin signal (shown in Fig. 1c) in primary human breast cancer. Patients are informed and asked for their participation through an approved ethics vote (mamma ethics vote No. 073-13-11032013).

The imaging of samples was conducted as follows: Primary breast cancer tumour samples were directly obtained from surgical procedures. Tumour tissues were sectioned into smaller pieces using sterile scalpels and subsequently fixed in 4% (wt/vol) paraformaldehyde in phosphate-buffered saline (PBS) for 30 minutes at room temperature, followed by multiple washes in PBS to remove residual fixative. Permeabilization was achieved by incubating samples in 1% Triton-X-100 overnight, with additional PBS washes afterwards. For staining, fixed samples were incubated overnight with 1 *µ*M Alexa Fluor 488-Phalloidin (Thermo Fisher Scientific, Waltham, MA, USA) to label actin structures, and 1 *µ*M SPY650-DNA (SC501, Spirochrome) to stain nuclear DNA. For imaging preparation and optical clearing, samples were mounted in ibidi IMM mounting medium (ibidi, Munich, Germany), with a refractive index (RI) of approximately 1.445. Imaging was conducted in *µ*-Slide 8Well chambers (ibidi), using a Zeiss LSM 980 confocal microscope equipped with AiryScan2. The Zeiss LD LCI Plan-Apochromat 25×/0.8 ImmKorr DIC M27 objective was used with a glycerol-water mixture (RI≈ 1.45) as the immersion medium to prevent refractive index mismatch between the sample and immersion medium. For optimal z-resolution, az-stack interval of 1 *µ*m was used.

### Proliferation fractions

A patient’s tumour tissue is stored in paraffin blocks from which sections (3-5 *µ*m) are cut and stained. Additional sections of the primary tumours were made approximately 10 *µ*m below the section plane of the analyzed HE-stained sections and stained with the mitotic marker Phospho-histone H3.

The quantitative analysis starts with a custom colour-deconvolution algorithm based on the work of Macenko et al. [31]. This operation separates the colour channels of the positive stain (brownish) and negative stain (blueish). Within the separate colour channels we employ the StarDist model [34] (pre-trained model for fluorescence images) to segment positive (mitotic activity) and negative (no mitotic activity) nuclei. The digital segments are filtered for areas *>* (1.5*µm*)^2^ · *π* and *<* (6*µm*)^2^ · *π* for reliable segments. The proliferation fraction *f*_*p*_ is then calculated as *f*_*p*_ = *N*_positive_*/*(*N*_positive_ + *N*_negative_) per patient.

### Mechanical probing of primary human breast cancer tissue

Primary human cancer samples taken during breast tumour surgeries were temporarily stored in a phosphate-buffered saline (PBS) solution at 4°C and measured within a few hours after removal. To prepare the tissue samples for AFM measurements, they were cut into thin slices of about 400*µ*m thickness using sharp scalpels and a McIlwain tissue chopper. The slices were then glued on a thin glass slide using

Histoacryl tissue adhesive (B.Braun Surgical, Rubí, Spain) and mounted in a 40-mm Petri dish (TPP, Trasadingen, Switzerland) filled with a few millilitres of modified ringer solution (B. Brown, Melsungen, Germany) to protect them from drying out. All measurements were carried out in a liquid environment at room temperature. Samples were measured using a Zeiss upright AxioZoom.V16 fluorescence microscope (Carl Zeiss Microscopy GmbH, Jena, Germany) equipped with the NanoWizard 4 XP setup and a 300 *µ*m HybridStage (Bruker Nano GmbH, Berlin, Germany). A modified Pointprobe-CONT cantilever (0.2 N/m) (Nanoworld AG, Neuchâtel, Switzerland) with an attached polystyrene bead (6 *µ*m diameter) was used as a spherical indenter for an increased contact area. Several thousand force indentation curves were measured across each sample surface with a set force of 2nN, covering an area of about 600*µ*m× 600*µ*m. For each force curve, the cantilever was held at constant height as soon as the set force was reached and sinusoidal oscillations were performed in a range from 10Hz-100Hz. After the mechanical probing was completed, Hoechst 33258 nucleic acid dye (Thermo Fisher Scientific, Waltham, USA) was added to the dish (mixing ratio 1:1000) to stain for DNA visualizing the nuclei positions to use for further tissue classification. The camera and cantilever positions were calibrated and after ∼30 min of incubation, the samples were imaged in tiles. Young’s moduli were obtained from a standard Hertz model fit [35, 36] with a Poisson’s ratio of 0.5 using the JPK data processing software V7.1.18 (Bruker Nano GmbH, Berlin, Germany). The viscoelastic properties, i.e. shear storage and shear loss, were calculated from the vibration data using a Sneddon model fit [36]. Self-written MATLAB scripts (MathWorks, Natick, USA) were then used to statistically analyze the data and to stitch the individual image tiles.

### Fitting Cluster Sizes and Shapes

Cancer cell clusters in the histological images are identified via an algorithm described in our previous work [3]. Subsequently, MATLAB’s *regionprops* function is used to measure areas and perimeters. We filter empirical cluster areas below (10*µm*)^2 ·^ *π*≈ 314*µm*^2^ since below this area it is not clear if the cluster contains more than one cell. Also, areas above 1.153· 10^5^*µm*^2^ are ignored due to data fluctuations. In the simulations, we identified individual clusters by using the inbuilt contour plot function in MATLAB with a single intermediate level. We then calculated the area and perimeter of the clusters from the contour matrix data, using the inbuilt polygon area and perimeter functions in MATLAB. The width of cluster boundaries (where 0.1 ≤*ϕ* ≤0.9) is approximately 5 LB = 25*µ*m, and we filter clusters smaller than this when extracting the power law exponents.

### Survival analyses, prognostic observables

We divide the patient collective presented in SI Table 1 into a random training set (30%; SI Table 2) and test set (70%; SI Table 2). As follow-up events, we take the *death from any cause* and *distant metastasis*. We censor all patients above 15 years of follow-up time to avoid events not related to cancer. Disease-free survival probabilities are estimated via the Kaplan-Meier estimator 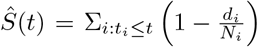,where *t*_*i*_ are time steps where at least one event happened, *d*_*i*_ is the number of events at *t*_*i*_ and *N*_*i*_ is the number of individuals known to have no event until time *t*_*i*_. This estimates the probability of not exhibiting certain events over time. For risk group separation we search for the maximum separation in the training set based on the log-rank statistic [37], 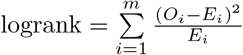, where *m* is the number of groups, *O*_*i*_ the total number of observed events and *E*_*i*_ the total number of expected events. Additionally, each risk group must contain at least 15% of patients in the training set to avoid unreliably small patient groups. The threshold found is then applied to the test set.

**Table 1.** Clinical characteristics of complete patient collective (N = 2012).

| Characteristic | Value (Proportion) |
| --- | --- |
| **Age** |  |
| mean ± std [range] | 58.77 ± 11.74 [25, 98] |
| **Grade** |  |
| 1 | 463 (0.23) |
| 2 | 1072 (0.53) |
| 3 | 470 (0.23) |
| missing | 7 (0) |
| **Size [cm]** |  |
| mean ± std [range] | 1.95 ± 1.41 [0.2, 16] |
| missing | 4 (0) |
| **Nodal Status** |  |
| pN0 | 1470 (0.73) |
| pN+ | 520 (0.26) |
| missing | 22 (0.01) |
| **ER Status** |  |
| ER 0 | 215 (0.11) |
| ER 1 | 1791 (0.89) |
| missing | 6 (0) |
| **PR Status** |  |
| PR 0 | 402 (0.2) |
| PR 1 | 1599 (0.79) |
| missing | 11 (0.01) |
| **Chemo Status** |  |
| Chemo 0 | 949 (0.47) |
| Chemo 1 | 493 (0.25) |
| missing | 570 (0.28) |

**Table 2.** Clinical characteristics of training set sopulation (N = 600).

| Characteristic | Value (Proportion) |
| --- | --- |
| **Age** |  |
| mean ± std [range] | 57.78 ± 10.83 [25, 98] |
| **Grade** |  |
| 1 | 156 (0.26) |
| 2 | 323 (0.54) |
| 3 | 120 (0.2) |
| missing | 1 (0) |
| **Size [cm]** |  |
| mean ± std [range] | 1.87 ± 1.41 [0.2, 15] |
| missing | 1 (0) |
| **Nodal Status** |  |
| pN0 | 464 (0.77) |
| pN+ | 129 (0.22) |
| missing | 7 (0.01) |
| **ER Status** |  |
| ER 0 | 56 (0.09) |
| ER 1 | 543 (0.91) |
| missing | 1 (0) |
| **PR Status** |  |
| PR 0 | 112 (0.19) |
| PR 1 | 485 (0.81) |
| missing | 3 (0.01) |
| **Chemo Status** |  |
| Chemo 0 | 287 (0.48) |
| Chemo 1 | 135 (0.23) |
| missing | 178 (0.3) |

As observables we use the *average cluster shape index* ⟨Ω⟩ per patient and the *normalized maximum defect-cancer distances* max(Δ_*DC*_)*/ξ*_*ECM*_ per patient. The maximum distances max(Δ_DC_) are normalized with the effective length scale of the ECM of the respective patient *ξ*_ECM_. *ξ*_ECM_ is determined by considering the radially averaged spatial pixel autocorrelations *C*(*r*) of the binary ECM mask using the *Wiener-Khinchin Theorem* [38]. The length at *C* = 1*/e* is then identified as *ξ*_ECM_. This normalization is necessary since ECM length scales vary across patients and thus affect the distributions Δ_DC_. Thus, max(Δ_DC_)*/ξ*_ECM_ presents a dimensionless, standardized measure without confounding effects of varying ECM sizes and associated length scales.

The thresholds found for risk group separation shown in the main text are ⟨Ω⟩ = 1.62 and max(Δ_DC_)*/ξ*_ECM_ = 0.81.

### Computational Model

We distinguish the cancer clusters and ECM using a phase-field *ϕ*, where *ϕ* = 1 labels the cancer clusters, and *ϕ* = 0 labels the ECM. This phase field relaxes to the minimum of the Landau-Ginzburg free energy

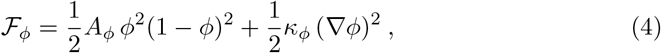

where *A*_*ϕ*_ and *κ*_*ϕ*_ are material parameters describing the surface tension between the species, and the sharpness of the interface. In this formalism, the surface tension between the two phases is 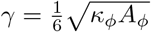 [39]. In the absence of growth, the dynamics of *ϕ* are given by the Cahn-Hilliard equation

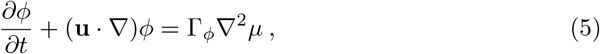

where **u** is the flow field discussed later, 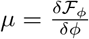 is the chemical potential, and Γ_*ϕ*_ is a mobility parameter.

We model the cancer clusters and ECM fibres inside each region using a nematic order tensor and a flow field, labelled by an index *i*. We choose *i* = 1 for the cancerous cells and *i* = 0 for the ECM. We define the concentration of phase *i* as *ϕ*^*i*^, where *ϕ*^1^ = *ϕ* and *ϕ*^0^ = 1 − *ϕ*. The nematic order of species *i* is described by the symmetric and traceless rank-2 tensor 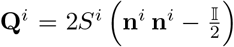, where **n**^*i*^ is the vector orientation field denoting the average orientation, and *S*^*i*^ is the magnitude of the nematic order parameter capturing the average degree of alignment. Each **Q**^**i**^ tensor relaxes to the minimum of the Landau de-Gennes free energy

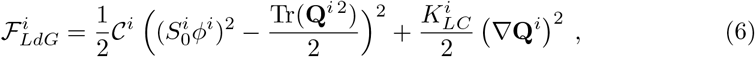

where, 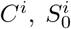 and 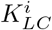 are material parameters representing the bulk free energy, average nematic order, and nematic elasticity respectively for each species. We define the molecular field

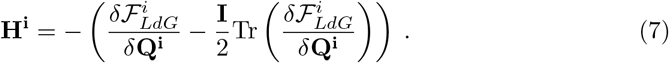

The flow field is modelled by a velocity field **u**, which satisfies the following equations:

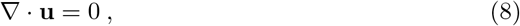

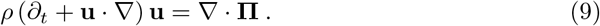

Here, *ρ* is the density of the fluid, and **Π**denotes the sum of all stresses acting on the fluid, composed of viscoelastic, active, nematic backflow, and capillary terms. The viscoelastic stress is modelled by the incompressible Kelvin-Voigt model [40–42]

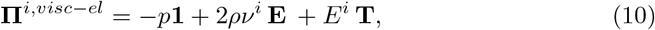

where *p* is an isotropic pressure term chosen to ensure incompressibility, ν^*i*^ is the kinematic viscosity, *E*^*i*^ is the elastic modulus, **E** = [(∇**u**) +(∇**u**)^*T*^]*/*2 is the strain rate tensor and **T** = ⎰ **E**(*t*) *dt* is the strain tensor. The total viscoelastic stress is

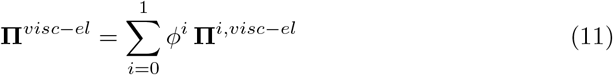

The nematic director field generates passive stresses upon stretching and distortion of the director field, called the nematic backflow. This is given by

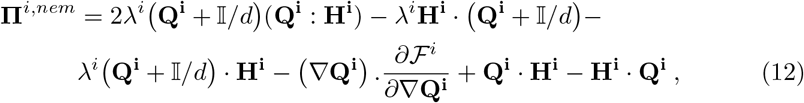

where *λ*^*i*^ is the flow-alignment parameter that describes the strength of alignment of the director axis with the strain rate in the fluid, and ℱ describes the total free energy.

Each species can produce active nematic stresses along the long axis of the director field, modelled by an active stress [8]

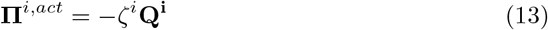

where ζ, the activity, characterizes the typical mechanical stresses exerted by each species on its environment. In our model, only the cancer clusters are active and hence ζ^0^ = 0.

The capillary stress models the stresses on the fluid across the boundary [43]. This is given by

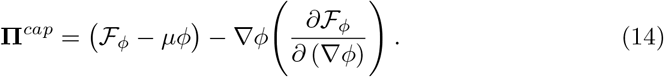

The total stress tensor acting on the fluid i s then given by

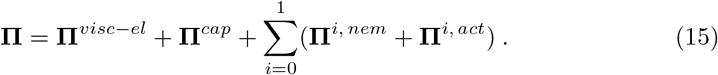

The dynamics of the director field is modelled by a Beris-Edwards equation [44]

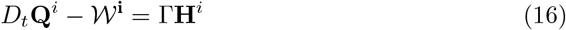

Here, the first term of the left-hand side is the convected time derivative, which describes the rate of change of **Q**^*i*^, which is being advected by the flow **u**. The second term *W*^*i*^ is the co-rotation term modelling the response of the director field to strain rate and vorticity in the flow. This is given by

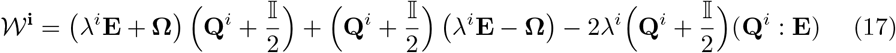

where **Ω** = [( ∇ **u**) − (∇ **u**)^*T*^]*/*2 is the vorticity tensor. The final term on the right-hand side of Eq. (16) is the molecular field (Eq. (7)), where Γ describes the rate of relaxation to the free-energy minimum.

### Simulation Parameters

We choose simulation parameters by first estimating the typical scales of the system.

#### Elastic and viscous moduli

We estimate the order of magnitude of the viscous *G*^*′′*^ [Pa] moduli within cancer clusters and ECM via frequency-dependent AFM measurements on a primary human breast cancer sample from the University Hospital in Leipzig. At low frequencies (*f* = 0.01Hz), the viscous moduli for both cancer tumours and ECM is around *G*^*′′*^∼ 10 *Pa*, resulting in an effective viscosity of *η*_1_ = *η*_2_ = 160Pa · s for the entire system. Cancer clusters fluidize the ECM in their vicinity [45], making it hard to accurately measure the elastic shear modulus in the system. Typical scales of the ECM shear elastic modulus can be estimated between 100 Pa - 1 kPa [13]. For a breast cancer sample measured with AFM, we estimate a shear modulus of the ECM ∼ 500 Pa (see Supplementary Note 1.2).

#### Lengths and velocities

Typical cell motility velocities are measured to be roughly *v*_0_ ∼3*µm/h* in vital primary human breast cancer [3]. Cancer cell tumours span a wide range of length scales, but typical cluster radii are on the scale of *R*∼ 10 −100*µm* (see size distribution in main text Fig. 3a).

#### Calibrating surface tension

We estimate this from cultured breast cancer cell spheroid fusion experiments reported in Ref. [11], which show that touching cancer clusters merge in *t* ∼ 36hr.

#### Active forces

Activity models the typical forces exerted by the cancer cells on their environment, which can include traction forces due to motility, and forces due to proliferation. The typical magnitude of these forces is around ζ∼400 Pa [15]. We caution that there will be considerable variation in these quantities between different biological samples.

With these scales, we can set normalized simulation units, hereafter referred to as ‘LB units’. We choose the typical length, time and stress units to be Δ*x* = 5 *µm*, Δ*t* = 30 *s*, and Σ = 5 kPa respectively. We use the elastic moduli *E*^0^ = 0.1 = 500 Pa (see Supplementary Note 1.2), *E*^1^ = 0.001 = 5 Pa, and activities in the range ζ^1^∈ [0.007, 0.2] = [35 Pa, 1000 Pa] (See Supplementary Note 1.3), and ζ^0^ = 0. We further use *ρ* = 40, ν^1^ = ν^0^ = 1, *A*_*ϕ*_ = *K*_*ϕ*_ = 1, and Γ_*ϕ*_ = 0.33 for the remaining fluid parameters.

The nematic parameters for the cancer clusters are chosen to be 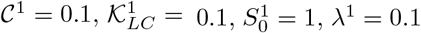 and Γ = 0.33. These are typical parameters for a nematic liquid crystal in an ordered phase, where the correlation length of the nematic orientation field is typically a few lattice spacings (∼ 10 − 20 *µm*).

For the ECM, we use 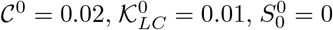,and *λ*^0^ = 0.1 for the nematic field instead. With these parameters, the Landau de-Gennes free energy has a weak effect on ECM dynamics (Eq.(16)), and topological defects and nematic alignment of the ECM are driven by active flows in the ECM.

With these parameter values, the typical active velocity is ∼6 ×10^−3^ (corresponding to 3.6 *µm hr*^−1^). Two droplets numerically initialized adjacent to each other merge, and the notch between two droplets of radii 20 (= 100*µm*) disappears in approximately 6 ×10^3^ timesteps (corresponding to ∼50 hr, see Supplementary Movie 4). More details about the parameter choice, and the robustness of the simulation results to changes in parameter values, are included in the Methods and Supplementary Note 1.3.

We ran simulations in a periodic box of size 400 ×400 for 400, 000 timesteps, with data recorded every 2000 timesteps. We either initialized the system with a circular droplet with radius 100, covering ∼20% of the simulation box (Movie 1), or with a uniform value of *ϕ* = 0.25 everywhere, resulting in many small droplets of *ϕ* = 1 covering 25% of the simulation domain (Movie 2). Both cases resulted in the same steady state. To obtain the results in Fig. 4, we ran simulations starting from a circular cluster of radius *R* = 40 in a periodic box of size 200× 200 for 84, 000 timesteps, with data collected every 2000 timesteps. In all cases, the nematic director field was initialized randomly and simulated for 100 pre-initialization steps without activity.

### Averaging over different activities

Different cancer clusters, within and across samples, show a spread of motility values [3, 11]. To compare simulation results to clinical data, we average over a distribution of activity values. The robustness of our results to different activity sampling schemes is discussed in Supplementary Note 1.3.

### Anchoring distributions

In the histology slides, we estimate boundary tangents from digitally interpolated (Matlabs in-built Savitzky-Golay-Filter *sgolayfilt*) pixel boundary segments. Angles between the respective tangents and ECM orientations in a square region (5*µm×* 5*µm*) in front of the boundary are calculated and the local median is used as the anchoring angle.

In the simulations, we calculate anchoring angles by measuring the angle between the ECM orientation and the maximum gradient of *ϕ* in regions close to the cancer-ECM boundary (0.05 ≤ *ϕ* ≤ 0.2).

### Decellularized 3D ECM imaging

In Supplementary Note 1.1.1, we present findings on the relationship between 2D and 3D nematic ordering within the context of 3D ECM. The imaging of samples was conducted as follows:

Samples of human liver-derived decellularized extracellular matrix (dECM) were thawed at +8°C and subsequently transferred to a 40 mm petri dish (TPP, Trasadingen, Switzerland) containing cooled phosphate buffered saline solution (PBS)(Life Technologies Limited, Paisley, United Kingdom). A major sample piece of dECM was cut using a scalpel such that a minor piece of ∼3× 3 mm size was obtained. A PBS solution containing 0.5% of Tetramethylrhodamine (TAMRA) was prepared. The dECM slice was submerged into the solution and incubated for∼ 24 hours at 37°C. After the labeling was complete, the sample was washed with PBS in a total of 3 washing steps with a 5 minute halt in between. The sample was subsequently placed in a *µ*-slide 18 well plate (Ibidi, Gräfelfing, Germany) and submerged in 500 *µ*l glycerol. For the microscopy, an Axio observer Z1/7 (Zeiss, Jena, Germany) equipped with a Yokogawa CSU-X1A 5000 spinning disk confocal scanning unit was used in fluorescence mode. The objective in use was an LCI Plan-Neo-fluar 25x/08 Imm (Zeiss, Jena, Germany). The staining excitation at 500 nm wavelength was performed using a CSU-X1 laser at 27% intensity. The image capturing was performed using a Hamamatsu Orca flash 4.0. The *µ*-slide containing the sample was placed on the holder stage and the objective was slowly approached to the sample until the glycerol droplet connected both the objective and sample vessel. We used the ZEN 2.6 (Zeiss, Jena, Germany) built-in image tessellation application and tessellated the sample into 8 equal-sized areas. The upper and lower z-boundary of the sample within the tessellated area was estimated by changing the focus. We divided the range from the lower to the upper boundary (∼405 *µ*m) yielding a total of 196 images with a step size of 2*µ*m in z-direction. The total recorded volume of 811 *µ*m x 1070 *µ*m x 405 *µ*m was processed by using ZEN 2.6 built-in image convolution and background subtraction algorithm.

## Acknowledgements

We thank Axel Niendorf, head of Pathology Hamburg-West, for months of valuable discussions, the provision of the histological database and the corresponding server structure. P.G. acknowledges doctoral scholarship funding from the University of Leipzig. S.B. acknowledges funding from the Crewe Graduate Award (Lincoln College, Oxford) and the Rudolf Peierls Centre for Theoretical Physics. J.M.Y. acknowledges support from ERC Advanced Grant Act-Bio (funded as UKRI Frontier Research Grant EP/Y033981/1).

## Author contributions

J.M.Y., J.A.K, S.B. and P.G. wrote the manuscript. J.M.Y., J.A.K, S.B. and P.G. conceptualized the study. S.B. and J.M.Y performed computer simulations, data analyses and data visualization. P.G. and K.L. performed quantitative analyses of histology slides. P.F. performed AFM measurements; P.F. and P.G. analysed the respective data. H.K. assisted data analysis, data interpretation and data visualization. I.S., A.D., J.W. prepared and provided 3D dECM. K.R. and S.R.M. imaged 3D dECM. P.G. analysed dECM. B.A. performed surgery and provided 3D breast cancer tissue. M.M. prepared and imaged 3D primary breast cancer. P.G. analysed 3D primary breast cancer. A-S.W. discussed histology data and helped conceptualize the study. J.M.Y. and J.A.K. supervised the study. All authors provided feedback and gave approval for the final version of the manuscript.

## Supplemental Notes

### S1.1 Dimensionality analyses

#### S1.1.1 ECM nematic order dimensionality

Histological H&E-stained slides are thin sections from 3D tumour tissue where the ECM is stained by eosin. Histological slides with eosin-stained ECM are prepared as the standard for all operable tumours. The sections are 3− 5*µm* in thickness. The measurable 2D orientation field arises from projecting the eosin-absorbance onto the plane visible by light microscopy. Therefore, as long as the nematic orientation is not perfectly perpendicular to the section, 2D order parameters should also indicate the local 3D order. We tested this with decellularized ECM (dECM) of human liver tissue stained for TAMRA (Fig. S1) and 3D spinning disk microscopy (see Methods). The dECM was provided by the Charité in Berlin (Ethics approval from the Ethikkommission der Charité – Universitätsmedizin Berlin exist for the use of tissue removed in the course of partial liver resection to obtain and cultivate liver cells and matrix components (EA1/289/16)). For comparison, we calculate the 2D scalar order parameter as described in the methods section in the main text for every *xy*-plane. 3D scalar order parameters are similarly calculated by considering the respective Q-tensor 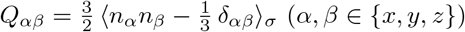 and diagonalizing it:

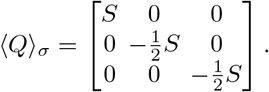

This enables us to compare *S*_2*D*_ and *S*_3*D*_ at every pixel of the digital volume of dECM for different coarse-graining length scales *σ*, see Fig. S1. As expected, we observe a good correlation between *S*_2*D*_ and *S*_3*D*_, with an average Pearson correlation coefficient of 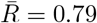.This is expected because when the section plane is not entirely orthogonal to the local nematic director, the projection onto the 2D plane yields information about the direction in the dimensions of the plane. However, if a nematic phase in 3D is completely orthogonal to the section, no information on the orientation can be obtained, which is why such cases are directly discarded by the constraints that orientation coherency must be *>* 0.1 and orientation energy *>* 0.01 during the structure tensor analysis (described in Methods). However, in real images, this is very rare.

#### S1.1.2 Cancer cell nematic order dimensionality

In the main text, we report 2D cell nematic ordering based on nucleus (DNA) intensities on histology slides. Here, we investigate the relation between 2D and 3D nematic cell ordering based on DNA (nucleus) and actin intensities in a 3D volume of primary human breast cancer shown in Fig. S2a, d and in the main text in Fig. 1c. Actin can be seen as a proxy of cancer cell outlines. Specimen and imaging details can be found in the Methods.

**Fig. S1.**
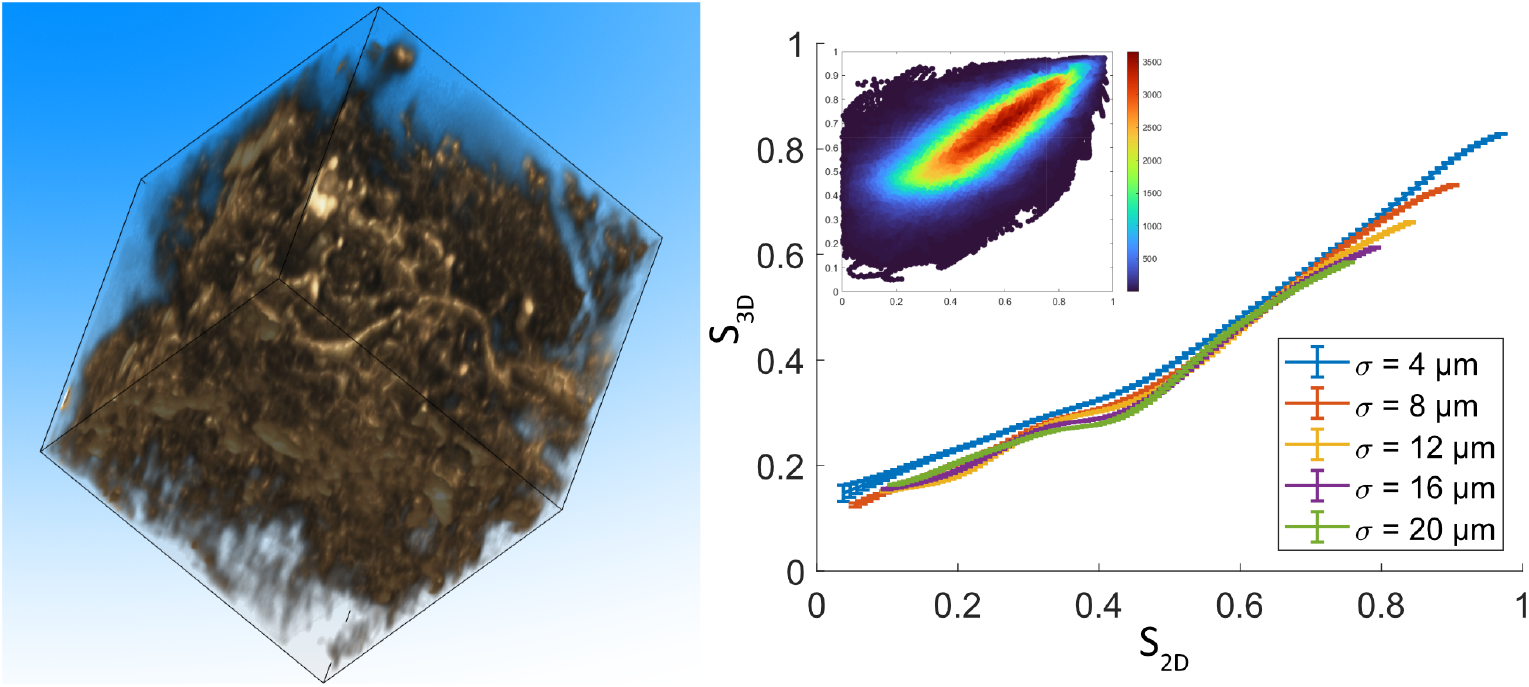
Relation between 2D and 3D ECM scalar nematic order parameters. Left: Digital dECM volume of TAMRA-stained primary human liver tissue. For visibility, low intensities are transparent. The edge length of the volume corresponds to 200*µ*m. Right: scalar nematic order parameter in 2D planes *S*_2*D*_ and in the volume *S*_3*D*_ for different coarse graining lengths *σ* of ⟨**Q⟩** _*σ*_. Data is averaged using a Gaussian moving average with a standard deviation of 0.05. The inset shows raw data for *σ* = 4*µm*. The average Pearson correlation coefficient across coarse graining length scales is *R* = 0.79.

We observe cell nematic ordering in 3D and 2D based on nuclei (Fig. S2b) and actin (Fig. S2e). For both cases, 2D and 3D nematic scalar order parameters are moderately correlated (Fig. S2c, f). We see that in 2D as well as 3D, actin and nucleus nematic order parameters are strongly correlated (Fig. S2g, h).

#### S1.1.3 Mapping sizes from three to two dimensions

The clinical images are two-dimensional projections of three-dimensional cancer clusters. However, the simulations consider two-dimensional clusters. The study of mapping from one to another is called stereology [S1]. An exact mapping can be derived for collections of perfect spheres [S2], and an analogous mapping can be derived for ellipsoids oriented isotropically [S3]. For more irregular shapes, studies show a strong correlation between the 2D observed size distributions and the real 3D volume distributions [S4].

As a specific example, we discuss the case of spherical clusters following the analysis in Ref. [S2] and show that the observed size distribution remains unchanged when looking at randomly chosen cross-sections. Consider a sphere or radius *R* and central cross-section area *A*_0_ = *πR*^2^. We consider a cross-section of the sphere corresponding to a plane at a distance *d* from its centre. The observed cross-sectional area is then *A* = *π*(*R*^2^ − *d*^2^), and hence 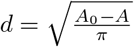.

We now assume the probability distribution of the *d* to be uniformly distributed across the radius of the sphere,

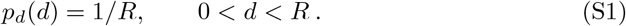

**Fig. S2.**
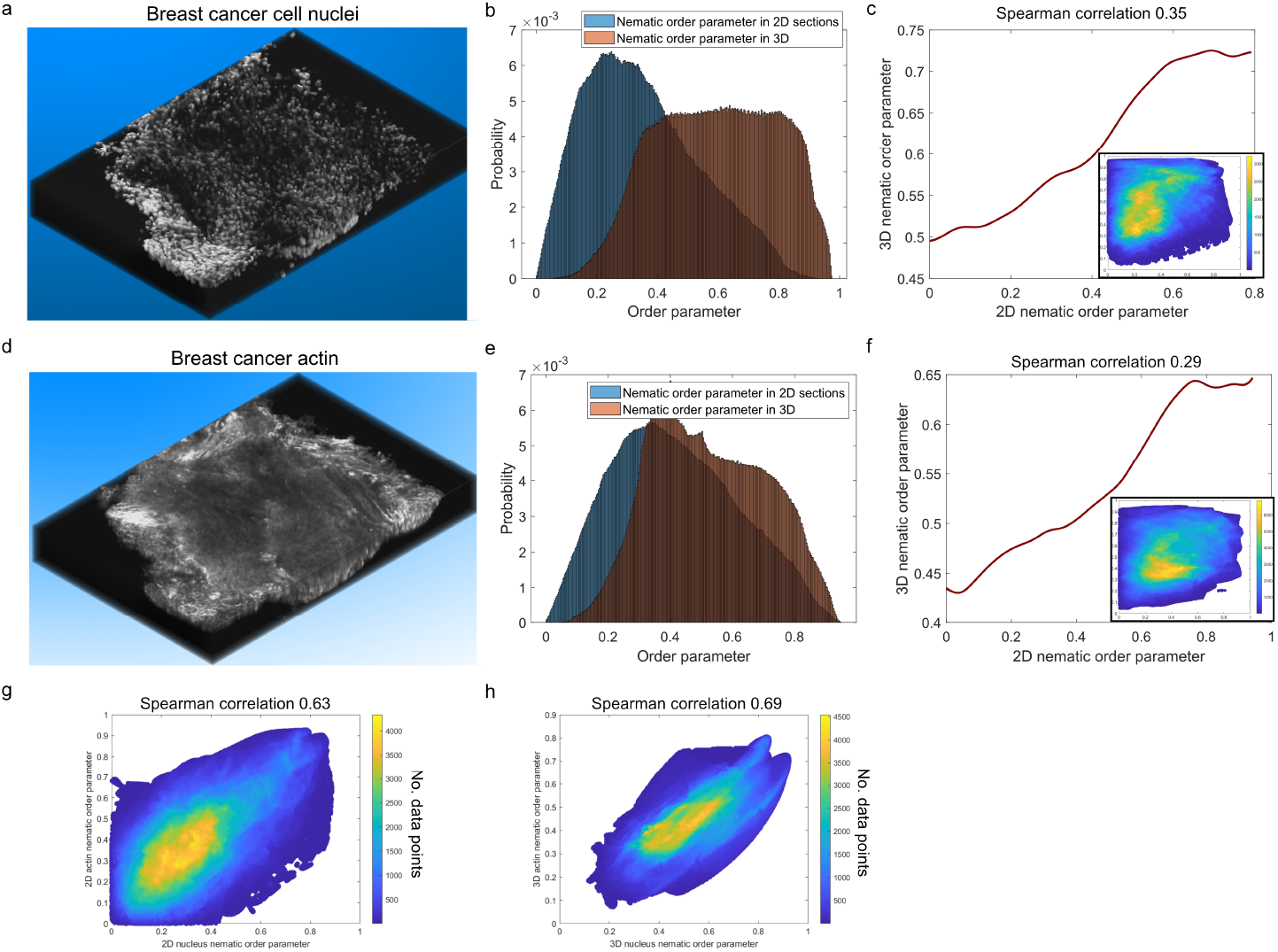
Relation between 2D and 3D cell scalar nematic order parameters in primary human breast cancer. **a** Volume of DNA intensities of primary human breast cancer (489*µm ×*721*µm ×*86*µm*). **b** Scalar nematic order parameter distributions based on nucleus (DNA) intensities in 2D and 3D. **c** Gaussian moving average (standard deviation 0.05) of the correlation between 2D and 3D nucleus nematic order parameters. Inset shows the raw data. Spearman correlation coefficient 0.35. **d** Volume of actin intensities of primary human breast cancer (489*µm* ×721*µm*× 86*µm*). **e** Scalar nematic order parameter distributions based on actin intensities in 2D and 3D. **f** Gaussian moving average (standard deviation 0.05) of the correlation between 2D and 3D actin nematic order parameters. Inset shows the raw data. Spearman correlation coefficient 0.29. **g** Correlation between 2D actin and nuclei nematic order parameters. Spearman correlation coefficient is 0.63. **h** Correlation between 3D actin and nuclei nematic order parameters. Spearman correlation coefficient is 0.69.

Changing variables to *A*, we find

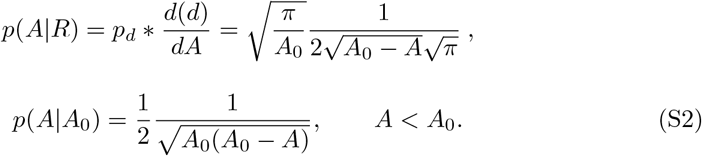

This is the probability of observing a cross-sectional area of *A*, for a sphere with a central cross-sectional area *A*_0_.

Finally, the observed distribution of *A*, for a distribution of sphere cross-sections *p*(*A*_0_), is given by

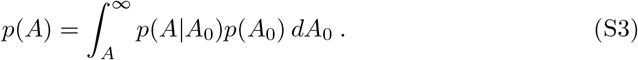

If *p*(*A*_0_) is distributed as a power law, Eq. (S3) implies that *p*(*A*) is distributed as a power law with the same exponent.

#### S1.1.4 Defect dimensionality

We suspect that 3D breast ECM contains disclinations whose crossing in the histological section planes can be identified by ± 1*/*2 defects.

To check the consistency of this, we ran simulations in 3D (active nematics comprising disclinations [S5]). For randomly chosen cross-sectional planes, we studied the director configurations where the 3D dislocations crossed the chosen plane. We expect from liquid crystal theory that the director configuration in the plane perpendicular to a dislocation line can vary between +1*/*2-like to −1*/*2-like, each occurring with equal probability, via twist configurations where the director field predominantly points normal to the plane [S5].

We observe that the in-plane defects (ignoring out-of-plane director components; Fig. S3b) indeed correspond to crossings of the disclinations with the plane (Fig. S3c). The number of defects measured in the plane is less than the number of dislocation crossings, as expected, as the twist configurations will not appear as 2D topological defects, and are not recorded in the 2D count. However, the estimated and real numbers exhibit a robust correlation with a Pearson correlation of *R* = 0.79 (Fig. S3d). Moreover, the number of +1*/*2 and −1*/*2 defects are, on average, equal (Fig. S3e) as expected, and as observed in the histological sections (Fig. S3f). These similarities strongly suggest that breast ECM contains 3D disclinations that can be recognized on 2D sections in the form of ± 1*/*2 defects

#### S1.1.5 Dimensionality of anchoring

In histological images of the ECM, we measure the 2D in-plane projection of the 3D orientation field. In this subsection, we estimate how projection affects the distribution of anchoring angles measured in Fig. 5 of the main text.

Consider a surface in the *y− z* plane, with its normal vector in the *x*-direction. We describe the 3D orientation of a nematic particle at the origin using a polar angle *θ*_3*D*_ and azimuthal angle φ_3*D*_, chosen such that *θ*_3*D*_ = 0^°^ corresponds to the *x*-axis. Thus, in Cartesian coordinates, the components of a normalized unit vector **n** in the (*θ*_3*D*_, φ_3*D*_) direction are given by *n*_*x*_ = cos *θ*_3*D*_, *n*_*y*_ = sin *θ*_3*D*_ cos φ_3*D*_, and *n*_*z*_ = sin *θ*_3*D*_ cos φ_3*D*_.

We further consider that we measure only the orientation field projected onto the *x*−*y* plane, losing information about the *z*-component. The measured orientation field in the plane is defined to be *θ*_2*D*_, such that components of a normalized unit vector **m** are given by *m*_*x*_ = cos *θ*_2*D*_, *m*_*y*_ = sin *θ*_2*D*_.

Note that *θ*_2*D*_ = 90^°^ corresponds to planar anchoring, and *θ*_2*D*_ = 0^°^ corresponds to homeotropic anchoring in 2D. In 3D, however, the ‘anchoring’ angle is the polar angle 4 *θ*_3*D*_ [S5]. Furthermore, the angles *θ*_3*D*_ and *θ*_2*D*_ are different, unless the 3D orientation field happens to lie in the *x*−*y* plane. Fig. S4a shows a schematic of the geometry under consideration. We study the mapping *θ*_3*D*_ →*θ*_2*D*_ by simulating nematics anchored perpendicular (*θ*_3*D*_ = 0^°^, shown in blue) or parallel (*θ*_3*D*_ = 90^°^, shown in red) to the surface, with different amplitudes of noise in the orientation angle (N=1000 samples for each noise level). In Fig. S4b, we find that the average anchoring angle measured in 2D is consistent with the average anchoring angle in 3D, showing that a preference for parallel anchoring in 2D corresponds to a preference for parallel anchoring in 3D. In Fig. S4c, we compare the fluctuations of the orientation field about the measured mean value. We find that the statistics of projected 2D anchoring are similar to the 3D anchoring in the parallel (planar) case, but the fluctuations of the projected anchoring angle are magnified in the case of perpendicular (homeotropic) anchoring.

**Fig. S3.**
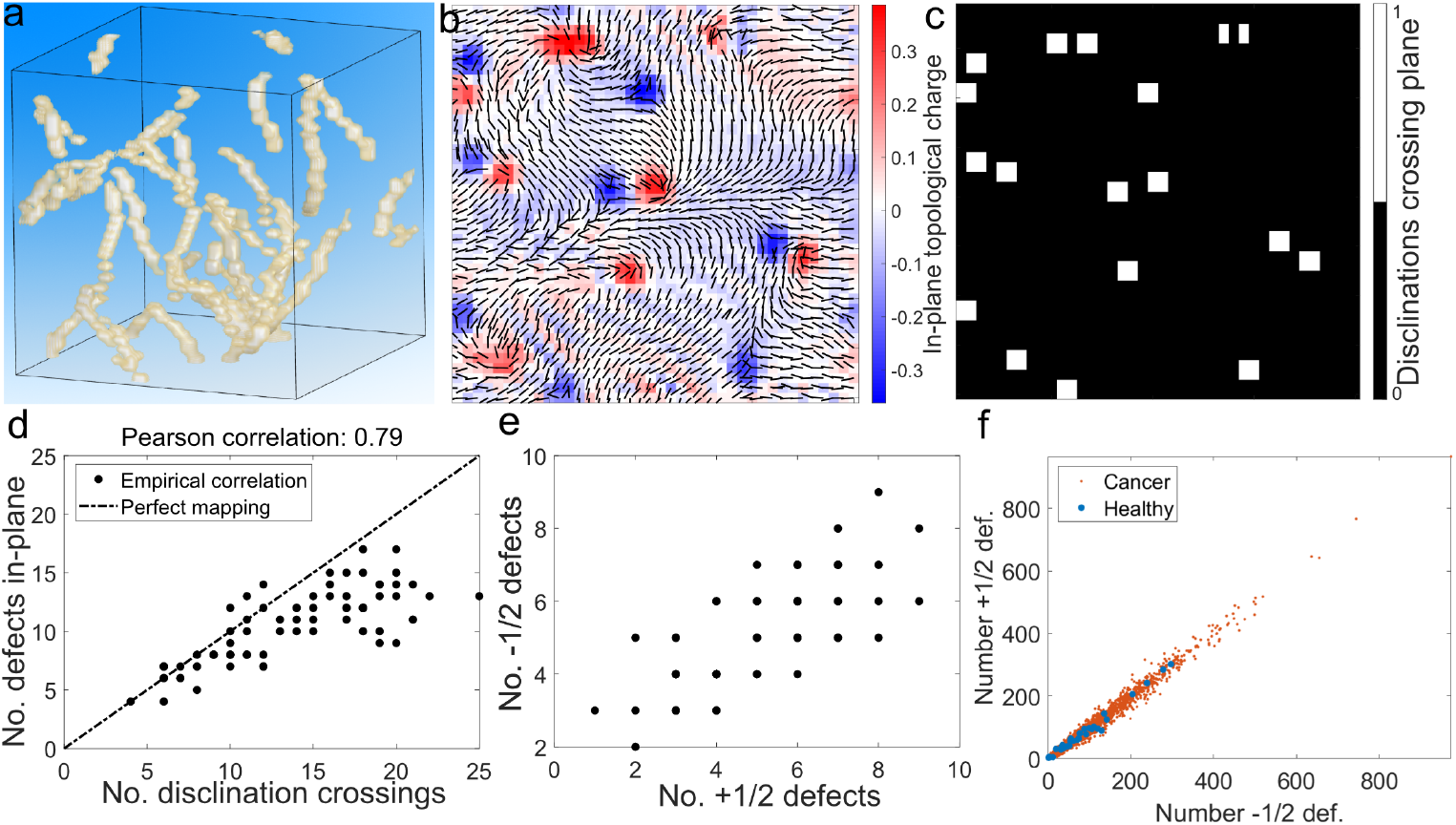
Analyses of simulation data of 3D director fields containing disclinations with respect to 2D sections. Half integer defects in sections indicate where disclinations are crossing the section. **a** Disclination lines in 3D. **b** A representative section with the normalized director field (*dx, dy*) containing no information about *dz*. The heatmap indicates the topological charge measured in-plane (see Methods). **c** Positions at which disclinations cross the plane shown in **b**, which correspond well with the defects identified from the 2D topological charge. **d** Number of ± 1*/*2 defects identified in-plane versus the actual number of disclination lines crossing the section. Defects identified in plane and dislocation crossings exhibit a robust correlation of *R* = 0.79. **e** Linear correlation between +1*/*2 and −1*/*2 defects measured in-plane, comparable to the linear scaling in the histological sections that is shown in **f. f** Linear correlation between number of +1*/*2 and −1*/*2 defects (*σ* = 5*µ*m) in the histological samples. Each dot corresponds to the respective numbers for one patient.

In the main text, we report the anchoring angle *θ*_*a*_. We define this as the angle subtended by the orientation axis of the projected ECM fibre to the cancer cluster tangent vector. Thus, *θ*_*a*_ = |90^°^ − *θ*_2*D*_|.

**Fig. S4.**
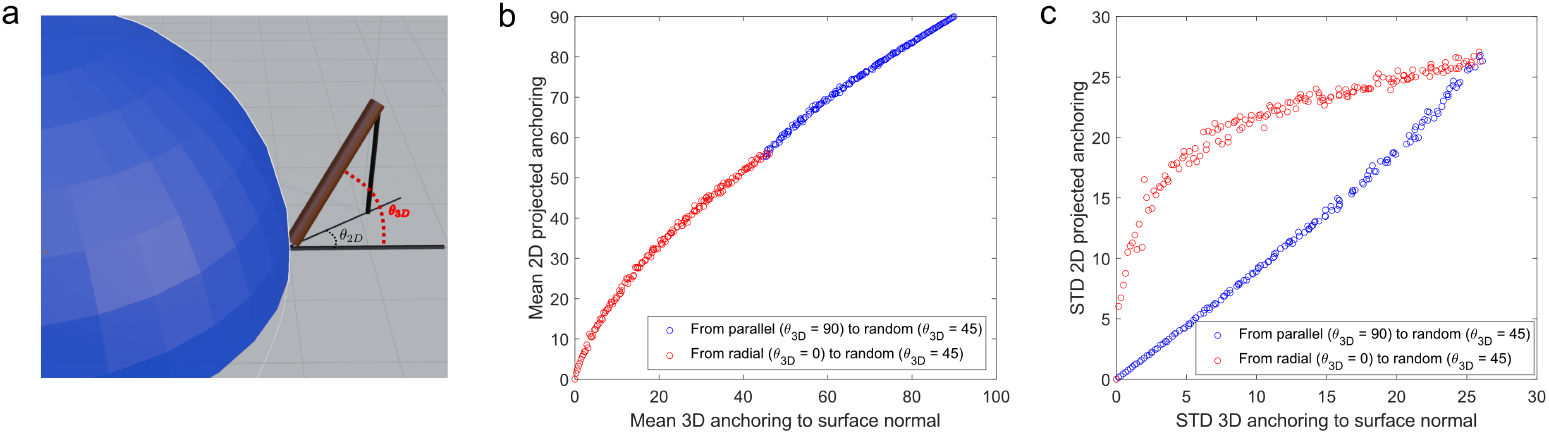
Relation between 3D anchoring and projected 2D anchorings. **a** Schematic of the 3D and 2D anchoring angles. **b** Mapping of the average anchoring angles in 3D to projected 2D average anchoring angles. **c** Mapping of standard deviations of anchoring angles in 3D to projected 2D anchoring angles.

### S1.2 Atomic force microscopy on primary breast cancer tissue

We performed AFM experiments on a primary breast cancer specimen after surgery to obtain order of magnitude elastic and viscous stress scales of cancer clusters and ECM (see Methods). With aid from pathologists, we roughly annotated the DNA image that is aligned to the mechanical measurements into dense cancer clusters and ECM. We group the mechanical data obtained from the AFM measurements into these regions. We then look at the frequency scans of the shear storage and shear loss moduli 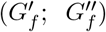 which we fitted by the fractional derivative Kelvin-Voigt model [S6] to extrapolate the low frequency regime of *f* ∼ 0.01 Hz relevant to the timescale of cancer activity (motility). For the ECM we estimate 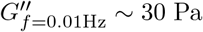 and 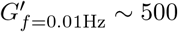 PA and for dense cancer cluster: 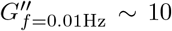 PA and 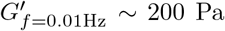 (Fig. S5a).

**Fig. S5.**
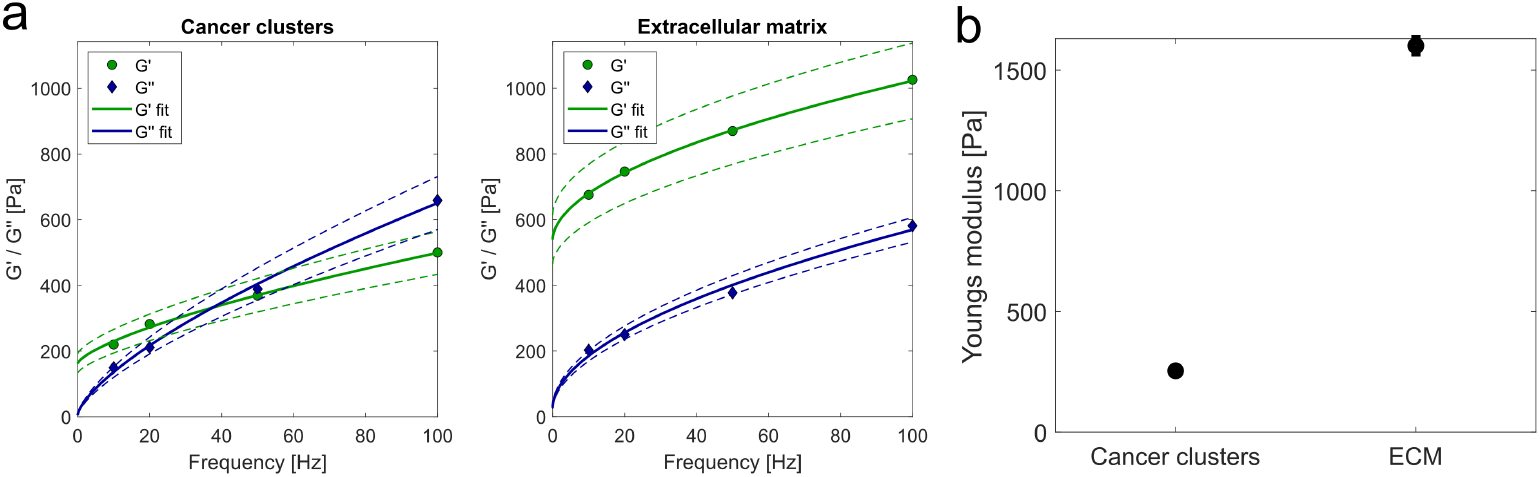
**a** Frequency-dependent AFM measurements of a primary human breast cancer sample (internal ID MCA 239) for different tissue regions: Cancer clusters, where DNA signal is very dense. Extracellular matrix: where no DNA signal is found. Frequency scans of the shear storage and shear loss 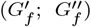 were fitted by the fractional derivative Kelvin-Voigt model [S6] to extrapolate the low frequency regime of relevance *f* = 0.01Hz. For the ECM we estimate 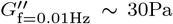 and 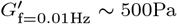 and for dense cancer cluster: 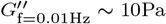 and 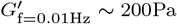. **b** Measured Young’s moduli in the different regions.

We also measured the Young’s modulus *Y* in the respective regions (Fig. S5b) to approximate the frequency-independent shear modulus of the ECM for the simulations. We observe that ⟨*Y*⟩ ∼ 1600 Pa. Since the measured Young’s modulus estimates the time-independent material elasticity when all relaxations have already taken place, we can estimate the respective shear modulus *G* via:

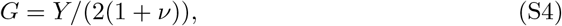

where ν is the Poisson ratio, which is around ν *≈*0.5 for biological materials [S7–S9]. Thus, we arrive at a rough estimate of the shear modulus of the ECM: *G*∼ 500 Pa which we use for our model.

### S1.3 Robustness of size and shape distributions to changing simulation parameters

When studying the distribution of cancer cluster sizes, we need to account for the fact that the motility of cancer clusters may be different across different clusters and different patients. Therefore, we average simulation results over a distribution of activity to compare to the histological data. In Fig. S6a, we find that changing the exact form of the activity distribution does not make a qualitative difference in the distribution of size data, as long as the sampling includes clusters with intermediate activity. Different sampling schemes have different area fractions of cancer clusters with a given activity. These are labelled by different colours and the relative weights for different activity values are shown in Fig. S6b. Systems with higher activity have a significantly higher number of cancer clusters, which are smaller in size (see Fig. 2c in the main text, and also Fig. S6c). These form the large majority of the observed clusters, dominating the statistics of the overall distribution. In the results presented in the text, we have used ‘Scheme 1’, which samples a broad spread of activity in cancer clusters.

We also looked at the sensitivity of our results to the chosen parameters of the model. The effect of changing activity is already shown in Fig 2b of the main text. Fig. S7a shows a snapshot of a dynamical steady state with the default parameters used in the text (ζ= 0.12). Fig. S7b shows the dynamical steady state when the ECM is modelled as a viscous fluid instead of a Kelvin-Voigt fluid (*E*^0^ = 0), in which case the cancer clusters are more elongated. Fig. S7c shows the dynamical steady state with the surface tension parameters doubled (*A*_*ϕ*_ = *K*_*ϕ*_ = 2). In this case, we find that the average cancer clusters are less elongated. In Fig. S7d, we look at the default parameters but with a much smaller area fraction (10% cancer cluster coverage instead of 25%). In this case, the dynamical steady state looks similar to the default parameters, but the area covered by the cancer clusters is much smaller.

Although the sizes and shapes of cancer clusters change upon changing parameters in individual simulations, the results for the resulting size and shape distributions show very little difference after averaging over a range of activities (see Fig. S7e and f). This is because activity is a critical parameter in all these simulations, and averaging over activity scans over the range of shapes and sizes observed with different activities.

**Fig. S6.**
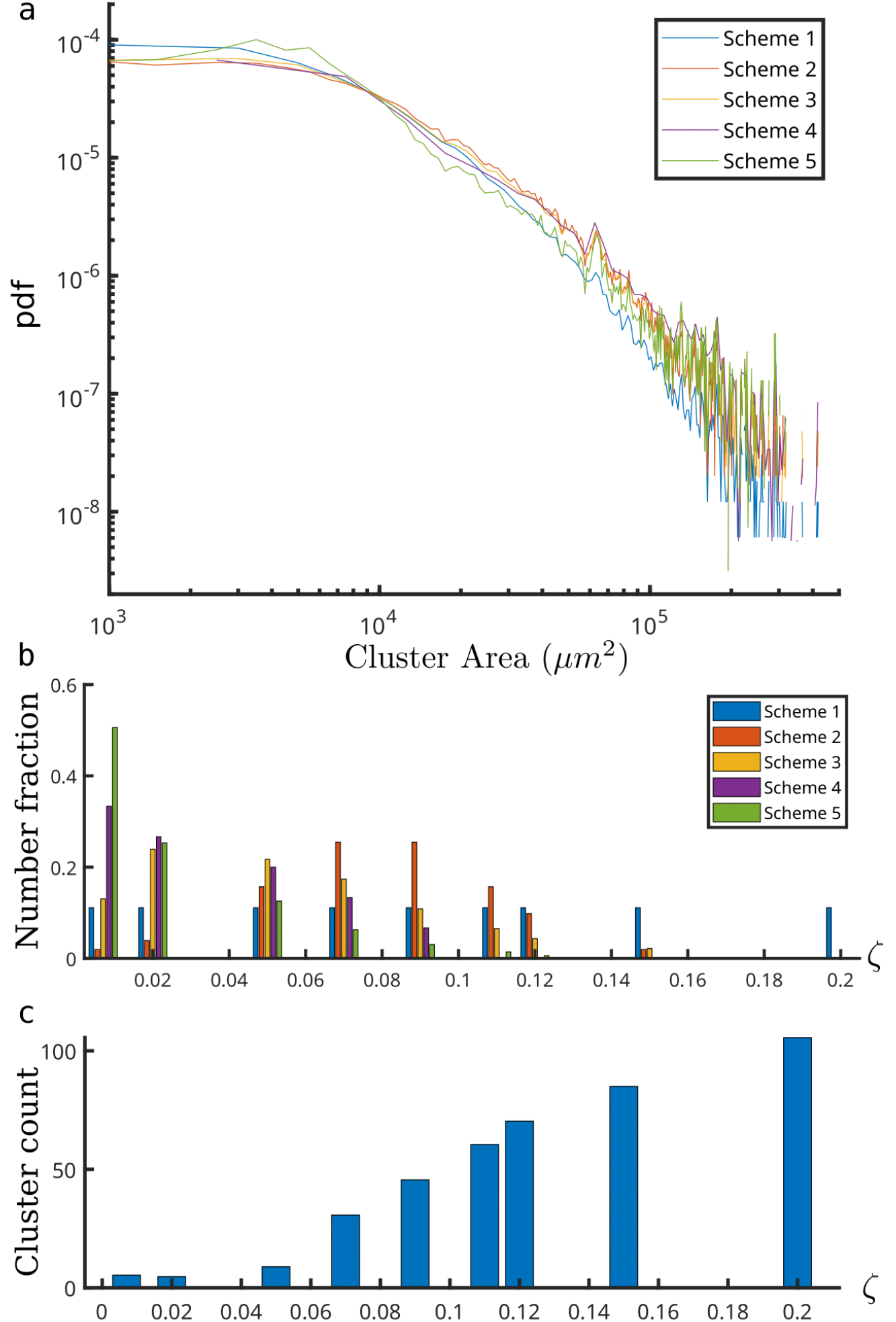
**a** Observed size distributions of cancer clusters for five different schemes of activity averaging, represented by different colours, all showing a similar power-law distribution. **b** Different activity averaging schemes, showing relative fractions of different activities in the system. **c** Bar plot showing the number of cancer clusters increases with activity, for a fixed area fraction of cancer cells.

**Fig. S7.**
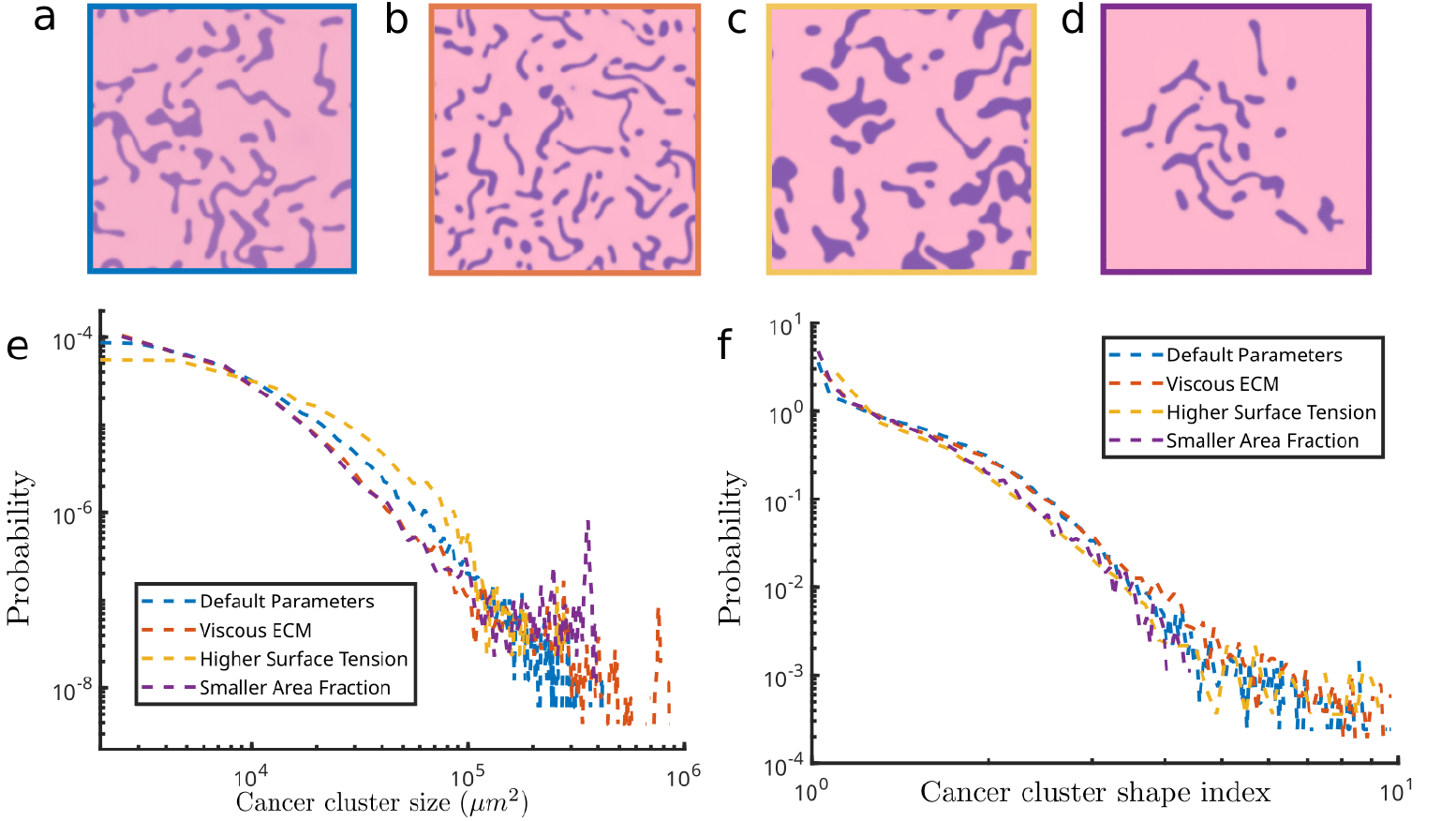
Robustness of cancer cluster sizes and shapes to simulation parameters. **a** Simulation snapshot with default simulation parameters. **b** Simulation snapshot with a viscous ECM. **c** Simulation snapshot for a higher surface tension. **d** Simulation snapshot for a smaller area fraction. **e** Distribution of size and **f** shape after activity averaging. The blue line shows results for parameters reported in the text, the red line shows results for purely viscous ECM, the yellow line shows results upon doubling the surface tension parameters, and the purple line shows a different area fraction.

### S1.4 Comparing healthy and cancer ECM

In Fig. S8b, we compare the scalar nematic order parameter between the ECM of healthy breast tissue and the ECM of invasive breast cancer for different coarse-graining lengths *σ*. We observed no statistical differences for *σ >* 7*µ*m. Below this threshold, healthy ECM exhibits slightly higher nematic order on small length scales and a higher topological defect density *ρ*_±1*/*2_ (Fig. S8c).

**Fig. S8.**
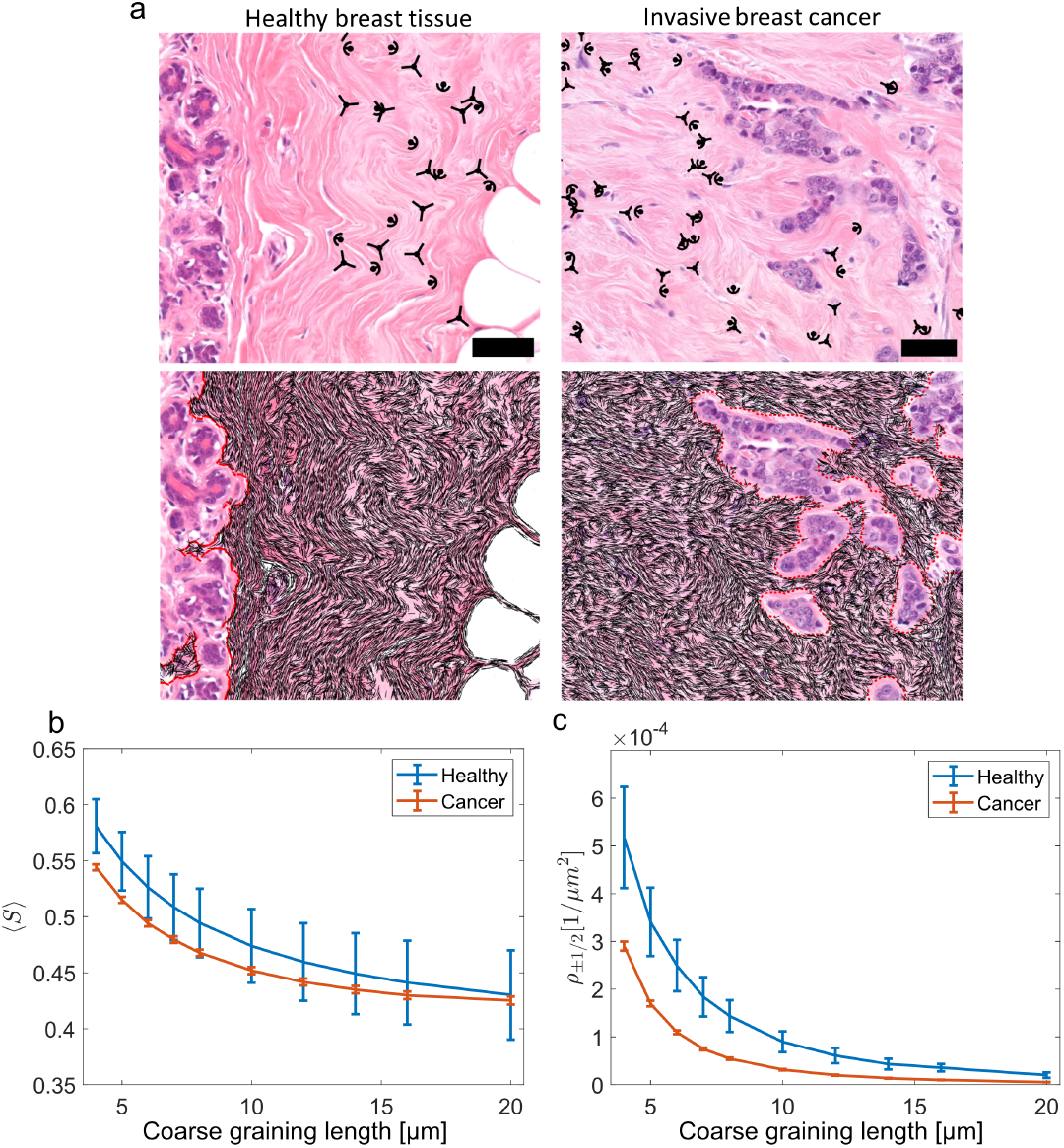
**a** Examples of histological sections of healthy breast tissue and invasive breast cancer tissue with topological defects and director fields overlaid. **b** Average nematic order ⟨*S*⟩ for different coarse-graining lengths *σ*. **b** Defect densities *ρ*_±1*/*2_ for different coarse-graining lengths *σ*. Defect densities are shown for healthy ECM (*N* = 32 patients) and invasive breast cancer ECM (*N* = 2012 patients). Number densities of ±1*/*2 defects scale with *ρ*_±1*/*2_ = *aσ*^−2^. Healthy: *a* = 0.0085 (*R*^2^ = 0.997); cancer: *a* = 0.0043 (*R*^2^ = 0.983).

### S1.5 Patient data

For our retrospective histopathological study, we investigated 2012 breast cancer patients. The analyses were based on H&E-stained digitized tissue sections of primary tumours. The cases considered were classified according to the WHO histological classification of breast carcinomas (3rd edition of 2003) which follows the TNM classification of the UICC (Union Internationale Contre le Cancer) in terms of grading and staging. All data used for this study comes from female patients. The clinic evaluated the sex of the patients by specification. The descriptive statistics of the patient collective are shown below.

In the following, the descriptive statistics are shown for the complete patient collective (Table 1), the training set (Table 2), and the test set (Table 3). For the defect-cancer distance observable patients without any defects in the ECM were neglected resulting in an overall 1982 patients for analysis.

**Table 3.** Clinical characteristics of test set population (N = 1412).

| Characteristic | Value (Proportion) |
| --- | --- |
| **Age** |  |
| mean ± std [range] | 59.19 ± 12.08 [25, 98] |
| **Grade** |  |
| 1 | 307 (0.22) |
| 2 | 749 (0.53) |
| 3 | 350 (0.25) |
| missing | 6 (0) |
| **Size [cm]** |  |
| mean ± std [range] | 1.99 ± 1.42 [0.4, 16] |
| missing | 3 (0) |
| **Nodal Status** |  |
| pN0 | 1006 (0.71) |
| pN+ | 391 (0.28) |
| missing | 15 (0.01) |
| **ER Status** |  |
| ER 0 | 159 (0.11) |
| ER 1 | 1248 (0.88) |
| missing | 5 (0) |
| **PR Status** |  |
| PR 0 | 290 (0.21) |
| PR 1 | 1114 (0.79) |
| missing | 8 (0.01) |
| **Chemo Status** |  |
| Chemo 0 | 662 (0.47) |
| Chemo 1 | 358 (0.25) |
| missing | 392 (0.28) |

Explanation of all of the characteristics:

- **Age**: Source: patient file. Explanation: Age at the time of surgery.
- **Grade**: Source: histological sample, defined by the examining pathologist following the Nottingham grading system in every breast cancer case. Explanation: Cancerous tissue can be morphologically assessed for the aggressiveness of the tumour based on the degree of differentiation (extent of degeneration and mitosis). Grading following the internationally accepted standard by UICC (Union Internationale Contre le Cancer): G1: well-differentiated G2: moderately differentiated G3: poorly differentiated
- **Size**: Source: histological slide, measured by the examining pathologist. Explanation: The tumour size affects prognosis as it is an indicator of tumour growth.
- **Lymph Node Status**: Source: histological preparation, measured by the examining pathologist. Explanation: Contains the information if any lymph nodes are invaded (Lymph Node Status = 1) or not (Lymph Node Status = 0). This is a strong and independent negative prognostic factor. When breast cancer begins to spread, the first metastatic site is often one of the neighbouring lymph nodes. From there, the tumour is able to spread further via the lymphatic system.
- **Hormone Receptor Status**: Source: Immunohistochemical staining, performed and analyzed by the pathologist. Explanation: The hormones estrogen and progesterone can affect the growth of breast cancer. They bind to hormone receptors of the cell, which then transmit the growth signal into the cell. To estimate whether a tumour grows hormone-dependent, the proportion of cells and the amount of corresponding hormone receptor-positive cells are analyzed. If more than one percent of all tumour cells are receptor-positive, it is assumed that the tumour is hormone-sensitive. This is indicated by ER+ (estrogen receptor-positive) and/or PR+ (progesterone receptor-positive).
- **Chemotherapy Status**: Source: Follow-up patient questionnaire: Information provided by the patient and/or treating physician. Explanation: Through chemotherapy, the cancer cells are attacked. The therapy is mainly used for tumours with aggressive growth. Before surgery, neoadjuvant chemotherapy can be used to shrink the size of the tumor.

## S2 Supplementary Videos

### S2.1 Video 1

Caption: Simulation data showing the dynamics of a single large active cancer droplet (dark pink) in a viscoelastic ECM (light pink). Active forces cause the cancer droplet to split apart into many tiny clusters, which form a dynamical steady state. In this state, individual cancer clusters move around, split and merge, however, the statistical properties of the clusters overall do not change with time.

### S2.2 Video 2

Caption: Simulation data showing the dynamics of many small active cancer droplets (dark pink) in a viscoelastic ECM (light pink). The droplets move and grow until they reach a dynamical steady state, identical to the one reached from a single large droplet initial condition. In this state, individual cancer clusters move around, split and merge, however, the statistical properties of the clusters overall do not change with time.

### S2.3 Videos 3a–e

Caption: Simulation data showing the dynamics of a single active cancer droplet (dark pink) in a viscoelastic ECM (light pink), with parameters corresponding to the representative points shown in Fig. 4 in the main text. All simulations are run from the same initial configuration.

### S2.4 Video 4

Caption: Simulation data showing the merging of two touching circular droplets of identical radius. The left subpanel shows the two clusters (deep pink) and the ECM (light pink) at any given time. The right subpanel plots the evolution of the aspect ratio of the bounding box as a function of time.

